# Evaluation and rational design of guide RNAs for efficient CRISPR/Cas9-mediated mutagenesis in *Ciona*

**DOI:** 10.1101/041632

**Authors:** Shashank Gandhi, Maximilian Haeussler, Florian Razy-Krajka, Lionel Christiaen, Alberto Stolfi

## Abstract

The CRISPR/Cas9 system has emerged as an important tool for various genome engineering applications. A current obstacle to high throughput applications of CRISPR/Cas9 is the imprecise prediction of highly active single guide. RNAs (sgRNAs). We previously implemented the CRISPR/Cas9 system to induce tissue-specific mutations in the tunicate *Ciona*. In the present study, we designed and tested 83 single guide RNA (sgRNA) vectors targeting 23 genes expressed in the cardiopharyngeal progenitors and surrounding tissues of *Ciona* embryo. Using high-throughput sequencing of mutagenized alleles, we identified guide sequences that correlate with sgRNA mutagenesis activity and used this information for the rational design of all possible sgRNAs targeting the *Ciona* transcriptome. We also describe a one-step cloning-free protocol for the assembly of sgRNA expression cassettes. These cassettes can be directly electroporated as unpurified PCR products into *Ciona* embryos for sgRNA expression *in vivo*, resulting in high frequency of CRISPR/Cas9-mediated mutagenesis in somatic cells of electroporated embryos.

We found a strong correlation between the frequency of an *Ebf* loss-of-function phenotype and the mutagenesis efficacies of individual *Ebf-*targeting sgRNAs tested using this method. We anticipate that our approach can be scaled up to systematically design and deliver highly efficient sgRNAs for the tissue-specific investigation of gene functions in *Ciona.*

## Introduction

A platform for targeted mutagenesis has been recently developed based on the prokaryotic immune response system known as CRISPR/Cas (Clustered Regularly Interspaced Short Palindromic Repeats/CRISPR-Associated) (Barrangou *et al.* 2007). In its most common derivation for genome engineering applications, the system makes use of a short RNA sequence, known as a single guide RNA (sgRNA) to direct the Cas9 nuclease of *Streptococcus pyogenes* to a specific target DNA sequence. (Jinek *et al.* 2012; Cong *et al.* 2013; Jinek *et al.* 2013; Mali *et al.* 2013). Although initial Cas9 binding requires a Protospacer Adjacent Motif (PAM) sequence, most commonly “NGG”, the high specificity of this system is accounted for by Watson-Crick base pairing between the 5’ end of the sgRNA and a 17-20bp “protospacer” sequence immediately adjacent to the PAM (Fu *et al.* 2014). Upon sgRNA-guided binding to the intended target, Cas9 generates a double stranded break (DSB) within the protospacer sequence. Imperfect repair of these DSBs by non-homologous end joining (NHEJ) often results in short insertions or deletions (indels) that may disrupt the function of the targeted sequence. Numerous reports have confirmed the high efficiency of CRISPR/Cas9 for genome editing purposes (Dickinson *et al.* 2013; Hwang *et al.* 2013; Wang *et al.* 2013a; Koike-Yusa *et al.* 2014; Paix *et al.* 2014; Shalem *et al.* 2014; Wang *et al.* 2014; Gantz and Bier 2015; Sanjana *et al.* 2016).

The tunicate *Ciona* is a model organism for the study of chordate-specific developmental processes (Satoh 2013). The CRISPR/Cas9 system was adapted to induce site-specific DSBs in the *Ciona* genome (Sasaki *et al.* 2014; Stolfi *et al.* 2014). Using electroporation to transiently transfect *Ciona* embryos with plasmids encoding CRISPR/Cas9 components, we were able to generate clonal populations of somatic cells carrying loss-of-function mutations of *Ebf*, a transcription-factor-coding gene required for muscle and neuron development, in F0-generation embryos (Stolfi *et al.* 2014). By using developmentally regulated *cis*-regulatory elements to drive Cas9 expression in specific cell lineages or tissue types, we were thus able to control the disruption of *Ebf* function with spatiotemporal precision. Following this proof-of-principle, tissue-specific CRISPR/Cas9 has rapidly propagated as a simple yet powerful tool to elucidate gene function in the *Ciona* embryo (Abdul-Wajid *et al.* 2015; Cota and Davidson 2015; Segade *et al.* 2016; Tolkin and Christiaen 2016).

We sought to expand the strategy to target more genes, with the ultimate goal of building a genome-wide library of sgRNAs for systematic genetic loss-of-function assays in *Ciona* embryos. However, not all sgRNAs drive robust CRISPR/Cas9-induced mutagenesis, and few guidelines exist for the rational design of highly active sgRNAs, which are critical for rapid gene disruption in F0. This variability and unpredictable efficacy demands experimental validation of each sgRNA tested. Individual studies have revealed certain nucleotide sequence features that correlate with high sgRNA expression and/or activity in CRISPR/Cas9-mediated DNA cleavage (Doench *et al.* 2014; Gagnon *et al.* 2014; Ren *et al.* 2014; Wang *et al.* 2014; Chari *et al.* 2015; Fusi *et al.* 2015; Housden *et al.* 2015; Moreno-Mateos *et al.* 2015; Wong *et al.* 2015; Xu *et al.* 2015; Doench *et al.* 2016). These studies have been performed in different organisms using a variety of sgRNA and Cas9 delivery methods and show varying ability to predict sgRNA activities across platforms (Haeussler *et al.* 2016).

Given the uncertainty regarding how sgRNA design principles gleaned from experiments in other species might be applicable to *Ciona*, we tested a collection of sgRNAs using our own modified tools for tissue-specific CRISPR/Cas9-mediated mutagenesis in *Ciona* embryos. We describe here the construction and validation of this collection using high-throughput sequencing of PCR-amplified target sequences. This dataset allowed us to develop a practical pipeline for optimal design and efficient assembly of sgRNA expression constructs for use in *Ciona*.

## Results

### High-Throughput sequencing to estimate sgRNA-specific mutagenesis rates

Previous studies using CRISPR/Cas9-based mutagenesis in *Ciona* revealed that different sgRNAs have varying ability to induce mutations (Sasaki *et al.* 2014; Stolfi *et al.* 2014). In order to test a larger number of sgRNAs and identify parameters that may influence mutagenesis efficacy, we constructed a library of 83 sgRNA expression plasmids targeting a set of 23 genes (**Table 1**). The majority of these genes are transcription factors and signaling molecules of potential interest in the study of cardiopharyngeal mesoderm development. The cardiopharyngeal mesoderm of *Ciona*, also known as the Trunk Ventral Cells (TVCs), are multipotent cells that invariantly give rise to the heart and pharyngeal muscles of the adult (Hirano and Nishida 1997; Stolfi *et al.* 2010; Razy-Krajka *et al.* 2014), thus sharing a common ontogenetic motif with the cardiopharyngeal mesoderm of vertebrates (Wang *et al.* 2013b; Diogo *et al.* 2015; Kaplan *et al.* 2015).

**Table 1.** Genes targeted for CRISPR/Cas9-mediated mutagenesis. The 23 genes targeted in the initial screen, each identified here by official gene symbol, aliases, and KyotoHoya identifier.

| # | Gene Symbol | Aliases | 2012 KyotoHoya ID |
| --- | --- | --- | --- |
| 1 | *Bmp2/4* | *Bone morphogenetic protein 2/4* | KH.C4.125 |
| 2 | *Ddr* | *Discoidin Domain Receptor Tyrosine Kinase 1/2* | KH.C9.371 |
| 3 | *Ebf* | *Collier/Olf/EBF; COE* | KH.L24.10 |
| 4 | *Eph.a* | *Ephrin type-A receptor.a; Eph1* | KH.C1.404 |
| 5 | *Ets.b* | *Ets/Pointed2* | KH.C11.10 |
| 6 | *Fgf4/6* | *Fibroblast growth factor 4/6; FGF, unassigned 1* | KH.C1.697 |
| 7 | *Fgf8/17/18* | *Fibroblast growth factor 8/17/18* | KH.C5.5 |
| 8 | *Fgfr* | *Fibroblast growth factor receptor* | KH.S742.2 |
| 9 | *Foxf* | *FoxF* | KH.C3.170 |
| 10 | *Foxg-r* | *Foxg-related; Orphan Fox-4; Ci-ZF248* | KH.C5.74 |
| 11 | *Fzd5/8* | *Frizzled5/8* | KH.C9.260 |
| 12 | *Gata4/5/6* | *GATA-a* | KH.L20.1 |
| 13 | *Hand* | *Heart And Neural Crest Derivatives Expressed 1/2* | KH.C14.604 |
| 14 | *Hand-r* | *Hand-related; Hand-like; NoTrlc* | KH.C1.1116 |
| 15 | *Htr7-r* | *5-Hydroxytryptamine Receptor 7-related* | KH.S555.1 |
| 16 | *Isl* | *Islet1/2* | KH.L152.2 |
| 17 | *Lef1* | *Lef/TCF* | KH.C6.71 |
| 18 | *Mrf* | *Muscle regulatory factor; MyoD* | KH.C14.307 |
| 19 | *Neurog* | *Neurogenin; Ngn* | KH.C6.129 |
| 20 | *Nk4* | *Nkx2-5; Tinman* | KH.C8.482 |
| 21 | *Rhod/f* | *RhoD/F; Rif* | KH.C1.129 |
| 22 | *Tle.b* | *Groucho2* | KH.L96.50 |
| 23 | *Tll* | *Tolloid-like; Tolloid* | KH.C12.156 |

We followed a high-throughput-sequencing-based approach to quantify the efficacy of each sgRNA, i.e. its ability to cause CRISPR/Cas9-induced mutations in targeted sequences in the genome (**Figure 1a-c**). The 83 sgRNA plasmids were co-electroporated with *Eef1a1>nls::Cas9::nls* plasmid. The ubiquitous *Eef1a1* promoter is active in all cell lineages of the embryo and has been used to express a variety of site-specific nucleases for targeted somatic knockouts in *Ciona* (Sasakura *et al.* 2010; Kawai *et al.* 2012; Sasaki *et al.* 2014; Stolfi *et al.* 2014; Treen *et al.* 2014). Each individual sgRNA + Cas9 vector combination was electroporated into pooled *Ciona* zygotes, which were then grown at 18°C for 16 hours post-fertilization (hpf; embryonic stage 25)(Hotta *et al.* 2007). Targeted sequences were individually PCR-amplified from each pool of embryos. Each target was also amplified from “negative control” embryos grown in parallel and electroporated with *Eef1a1>nls::Cas9::nls* and “*U6>Negative Control*” sgRNA vector. Agarose gel-selected and purified amplicons (varying from 108 to 350 bp in length) were pooled in a series of barcoded “targeted” and “negative control” Illumina sequencing libraries and sequenced using the Illumina MiSeq platform.

**Figure 1.**
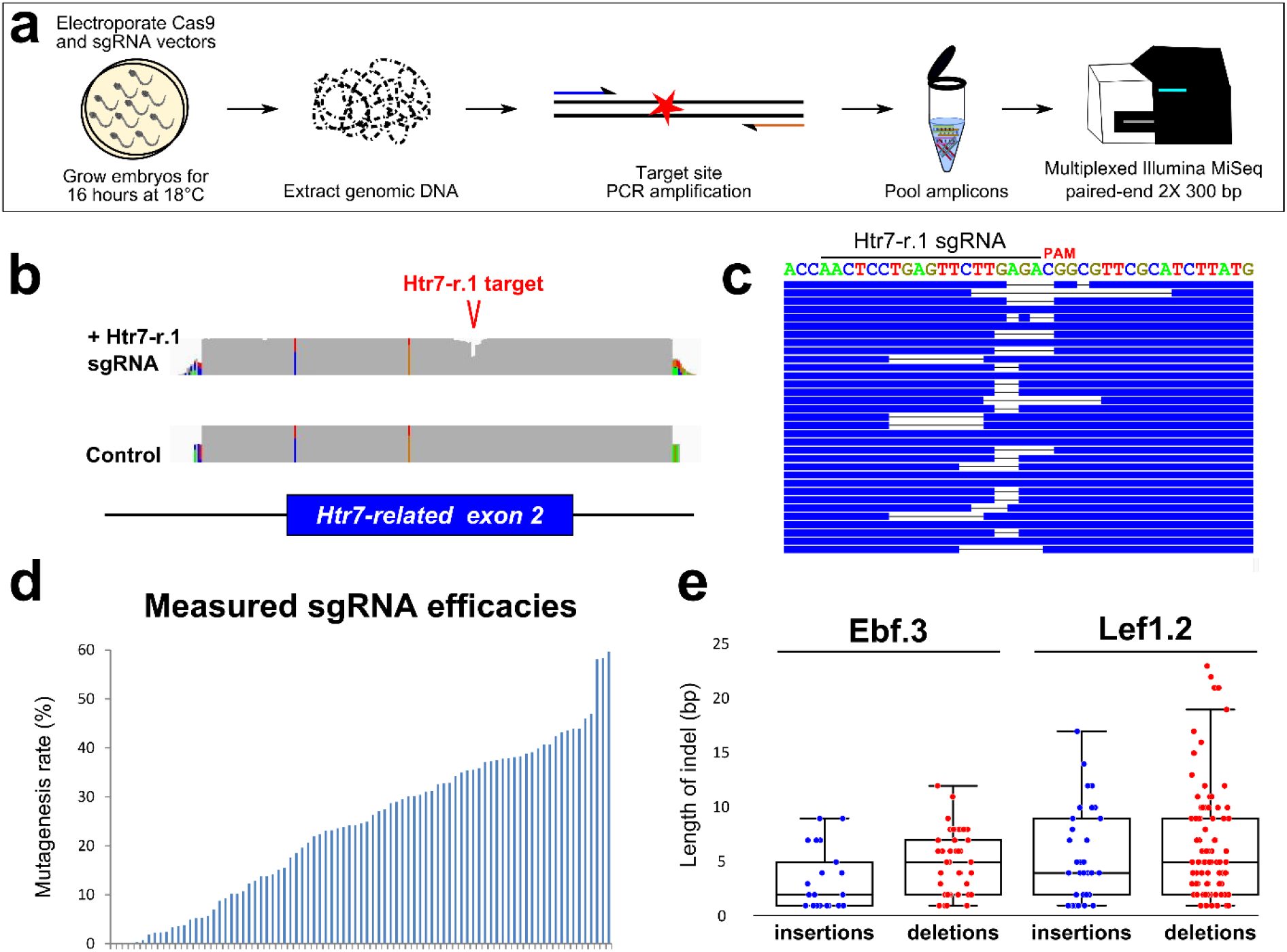
Next-Generation Sequencing approach to validating sgRNAs for use in Ciona embryos. **a)** Schematic for next-generation sequencing approach to measuring mutagenesis efficacies of sgRNAs expressed in F0 *Ciona* embryos. See results and materials and methods for details. **b)** Representative view in IGV browser of coverage (grey areas) of sequencing reads aligned to the reference genome. “Dip” in coverage of reads from embryos co-electroporated with *Eef1a1>Cas9* and *U6>Htr7-r.1* sgRNA vector indicates CRISPR/Cas9-induced indels around the sgRNA target site, in the 2nd exon of the *Htr7-related* (*Htr7-r*) gene. Colored bars in coverage indicate single-nucleotide polymorphisms/mismatches relative to reference genome. **c)** Diagram representing a stack of reads bearing indels of various types and sizes, aligned to exact target sequence of Htr7-r.1 sgRNA. **d)** Plot of mutagenesis efficacy rates measured for all sgRNAs, ordered from lowest (0%) to highest (59.63%). Each bar represents a single sgRNA. **e)** Box-and-whisker plots showing the size distribution of insertions and deletions caused by Ebf.3 or Lef1.2 sgRNAs.

Alignment of the resulting reads to the reference genome sequence (Satou *et al.* 2008) revealed that targeted sites were represented on average by 16,204 reads, with a median of 3,899 reads each (**Supplementary Table 1**). The ability of each sgRNA to guide Cas9 to induce DSBs at its intended target was detected by the presence of insertions and deletions (indels) within the targeted protospacer + PAM. The ratio of [indels]/[total reads] represents our estimation of the mutagenesis efficacy of the sgRNA (**Figure 1b-d, Supplementary Table 1**). For each mutation, an unknown number of cell divisions occur between the moment Cas9 induces a DSB and the time of sample collection and genomic DNA extraction. This prevents us from quantifying the true mutagenesis rates. However, we can surmise that this applies comparably to all sgRNAs, such that our values still represent an accurate ranking of mutagenesis efficacy.

For simplicity, we did not count single nucleotide point mutations, even though a fraction of them may result from NHEJ-repair of a DSB event. Our data indicated that all sgRNAs (with the exception of Neurog.2) were able to induce DSBs, with estimated efficacies varying from 0.05% (Ebf.4) to 59.63% (Htr7-r.2). Although each sgRNA was tested only once, we did not observe any evidence of electroporation variability or batch effects that may have confounded our estimates (**Supplementary Figure 1**).

This conservative approach most likely underestimates the actual mutagenesis rates. First, we excluded point mutations potentially resulting from imperfect DSB repair. Second, but more importantly, amplicons from transfected cells are always diluted by wild-type sequences from untransfected cells in the same sample, due to mosaic incorporation of sgRNA and Cas9 plasmids. Indeed, we previously observed an enrichment of mutated sequences amplified from reporter transgene-expressing cells isolated by magnetic-activated cell sorting (representing the transfected population of cells) relative to unsorted cells (representing mixed transfected and untransfected cells) (Stolfi *et al.* 2014). In that particular example, the estimated mutagenesis efficacy induced by the Ebf.3 sgRNA was 66% in sorted sample versus 45% in mixed sample. This suggests the actual efficacies of some sgRNAs may be up to 1.5-fold higher than their estimated rates.

Analysis of unique indels generated by the activity of two efficient sgRNAs, Ebf.3 and Lef1.2, indicated a bias towards deletions rather than insertions, at a ratio of roughly 2:1 deletions:insertions (**Figure 1e**). However, these two sgRNAs generated different distributions of indel lengths, indicating indel position and size may depend on locus-specific repair dynamics as previously shown (Bae *et al.* 2014).

Numerous studies have reported the potential off-target effects of CRISPR/Cas9 in different model systems (Fu *et al.* 2013; Hsu *et al.* 2013; Pattanayak *et al.* 2013; Cho *et al.* 2014). For this study, we were able to mostly select highly specific sgRNAs, owing to the low frequency of predicted off-target sequences in the small, A/T-rich *Ciona* genome (See **Discussion** for details). To test the assumption that off-target DSBs are unlikely at partial sgRNA seed-sequence matches, we analyzed the mutagenesis rates at two potential off-target sites that matched the protospacer at the 10 and 8 most PAM-proximal positions of the Ebf.3 and Fgf4/6.1 sgRNAs, respectively. We did not detect any mutations in 5,570 and 6,690 reads mapped to the two loci, respectively, suggesting high specificity of the sgRNA:Cas9 complex to induce DSBs only at sites of more extensive sequence match.

### Identifying sequence features correlated with sgRNA efficacy

We analyzed our dataset for potential correlations between target sequence composition and sgRNA-specific mutagenesis rate (excluding the Bmp2/4.1 sgRNA because only two reads mapped to its target sequence). We hypothesized that, if mutagenesis efficacy can be predicted by nucleotide composition at defined positions in the protospacer and flanking sequences, then comparing the target sequences of the most or least active sgRNAs in our dataset should reveal features that affect CRISPR/Cas9 efficacy in *Ciona*. To that effect, we performed nucleotide enrichment analyses for the top and bottom 25% sgRNAs ranked by measured mutagenesis efficacy (**Figure 2**)(Schneider and Stephens 1990; Crooks *et al.* 2004). For the top 25% sgRNAs, guanine was overrepresented in the PAM-proximal region, while the ambiguous nucleotide of the PAM (‘N’ in ‘NGG’) was enriched for cytosine. We also observed an overall depletion of thymine in the protospacer sequence for the top 25% sgRNAs, likely due to premature termination of PolIII-driven transcription as previously demonstrated (Wu *et al.* 2014). Among the bottom 25% sgRNAs, we observed a higher representation of cytosine at nucleotide 20 of the protospacer (**Figure 2**). All these observations are consistent with the inferences drawn from previous studies, suggesting that certain sgRNA and target sequence features that influence Cas9:sgRNA-mediated mutagenesis rates are consistent across different experimental systems (Gagnon *et al.* 2014; Ren *et al.* 2014; Wu *et al.* 2014; Chari *et al.* 2015; Moreno-Mateos *et al.* 2015; Haeussler *et al.* 2016).

**Figure 2.**
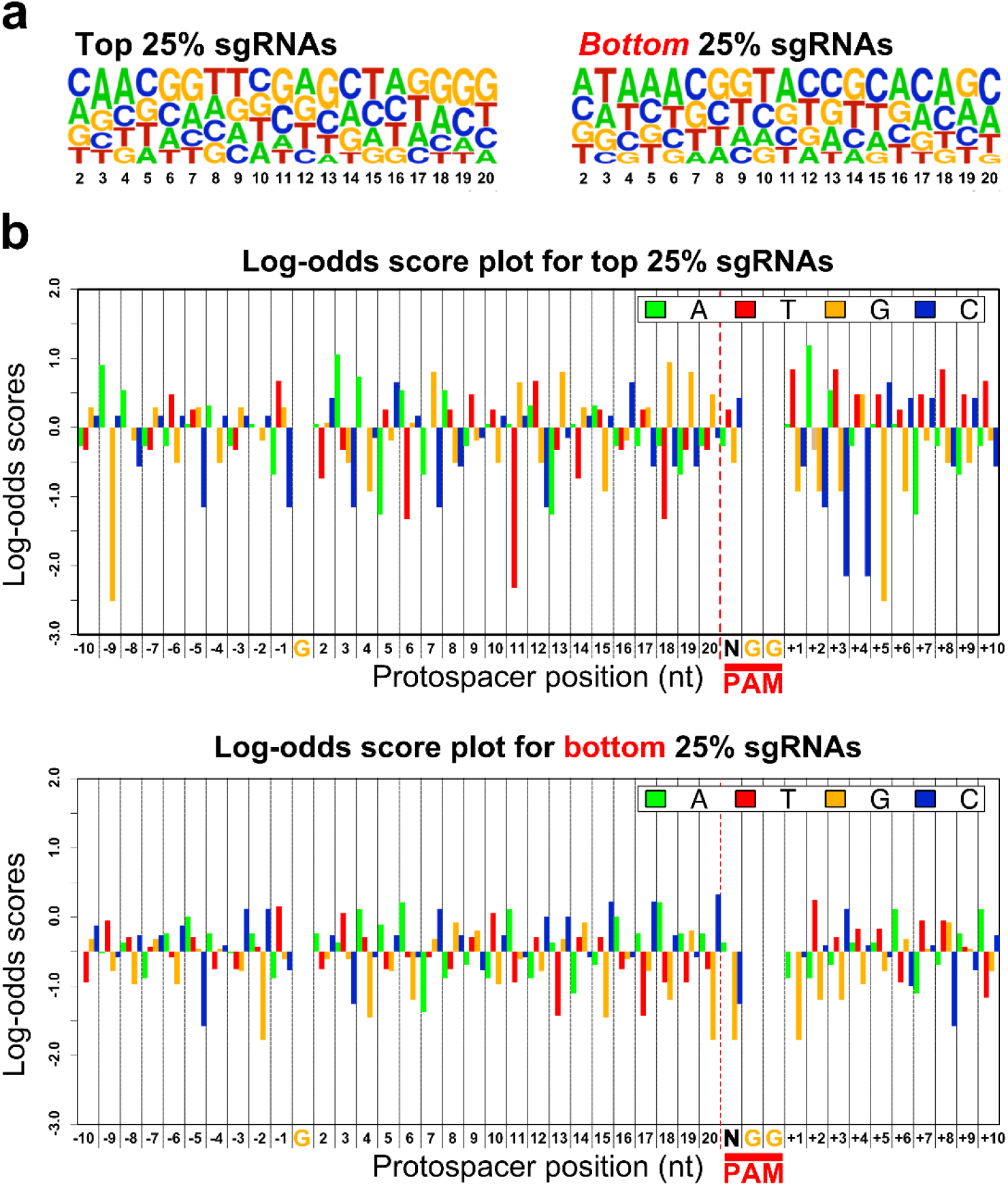
Correlations between sgRNA sequence composition and mutagenesis efficacy. **a)** WebLogos representing the nucleotide composition at each variable position of the protospacer (nt 2-20, X axis), in top 25% and bottom 25% performing sgRNAs. **b)** Log-odds scores depicting the frequency of occurrence for nucleotides in the top 25% and bottom 25% sgRNAs, at all positions of the protospacer, PAM, and flanking regions. Position “1” of the protospacer has been omitted from the analysis, due to this always being “G” for PolIII-dependent transcription of U6-promoter-based vectors. Likewise, the “GG” of the PAM has also been omitted, as this sequence is invariant in all targeted sites. Positive and negative log-odds scores reflect abundance and depletion, respectively, of a specific nucleotide at a given position relative to its random occurrence in our sample space. See **materials and methods**for details.

Several studies in different model organisms have similarly examined the nucleotide preferences amongst sgRNAs inducing high or low rates of mutagenesis, and put forth predictive heuristics and/or algorithms for the rational design of highly active sgRNAs (Doench *et al.* 2014; Gagnon *et al.* 2014; Ren *et al.* 2014; Wang *et al.* 2014; Chari *et al.* 2015; Fusi *et al.* 2015; Housden *et al.* 2015; Moreno-Mateos *et al.* 2015; Wong *et al.* 2015; Xu *et al.* 2015; Doench *et al.* 2016). These various algorithms have been evaluated, summarized, and aggregated by the CRISPR sgRNA design web-based platform CRISPOR (Haeussler *et al.* 2016). According to this meta-analysis, the “Fusi/Doench” algorithm (“Rule Set 2”)(Fusi *et al.* 2015; Doench *et al.* 2016) is the best at predicting the activities of sgRNA transcribed *in vivo* from a U6 small RNA promoter transcribed by RNA Polymerase III (PolIII), while CRISPRscan is recommended for sgRNAs transcribed *in vitro* from a T7 promoter (Moreno-Mateos *et al.* 2015).

*Ciona* rearing conditions differ from most other model systems, being a marine invertebrate that develops optimally at 16-24°C (Bellas *et al.* 2003). Moreover, most CRISPR/Cas9-based experiments in *Ciona* rely on *in vivo* transcription of sgRNAs built with a modified “F+E” backbone (Chen *et al.* 2013) by a *Ciona*-specific U6 promoter (Nishiyama and Fujiwara 2008). We used CRISPOR to calculate predictive scores for all sgRNAs in our data set and compare these scores to our mutagenesis efficacy measurements for each (**Figure 3a, Supplementary Table 1**). We found that indeed the Fusi/Doench score best correlated with our measured sgRNA efficacies, with a Spearman’s rank correlation coefficient (rho) of 0.435 (p=3.884e-05)(**Figure 3a**). CRISPRscan, the algorithm based on *in vitro*-, T7-transcribed sgRNAs injected into zebrafish, was less predictive (rho = 0.344, p=0.001435)(**Supplementary Table 1**), supporting the conclusion that sgRNA expression method (e.g. U6 vs. T7) accounts for an important parameter when choosing an sgRNA design algorithm. Scores computed by other published algorithms available on the CRISPOR platform did not yield good correlations with our measurements (**Supplementary Table 1**), indicating these are perhaps not suited for predicting sgRNA activity in *Ciona*.

**Figure 3.**
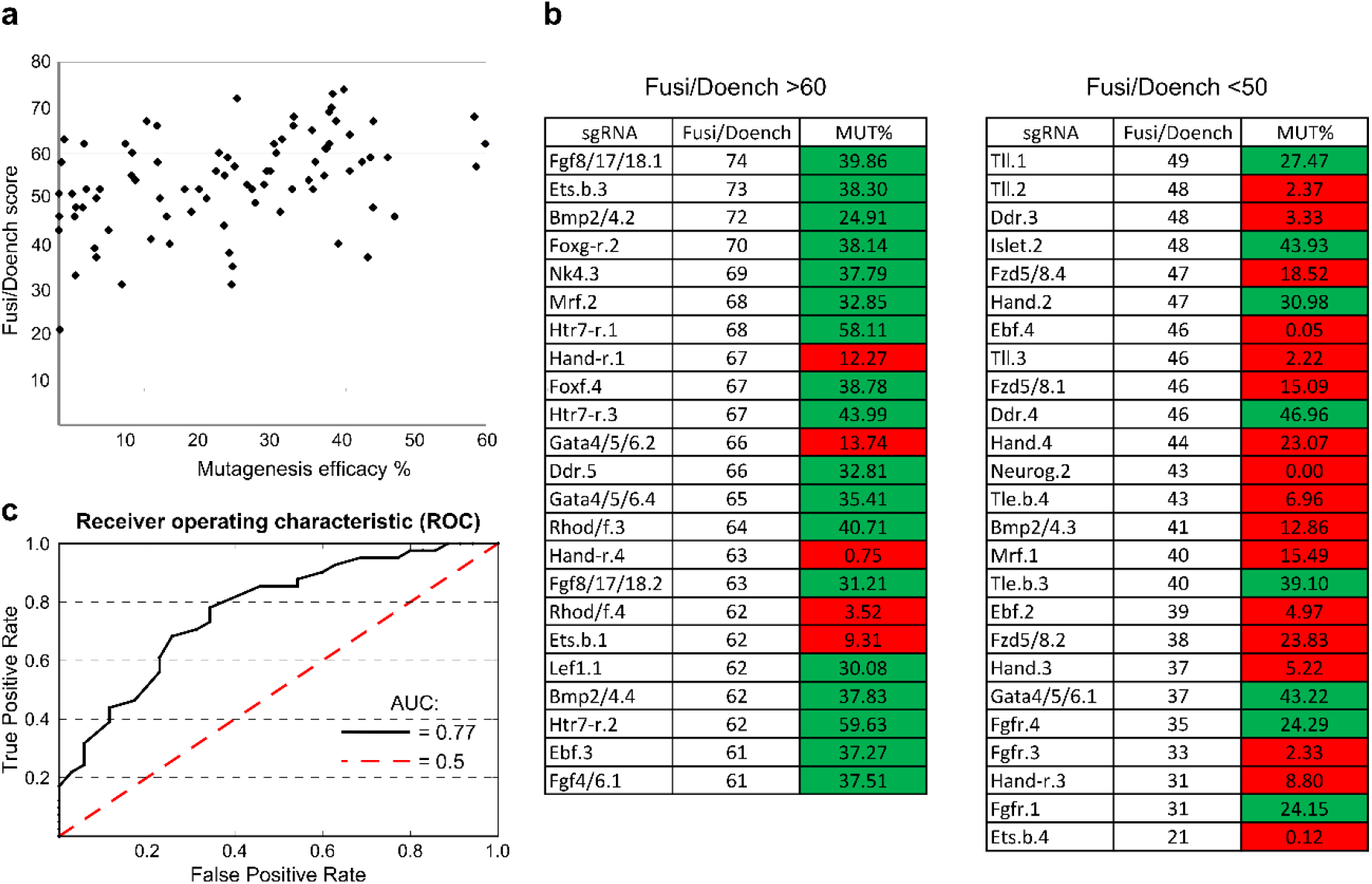
Using Fusi/Doench scores to predict mutagenesis efficacies of sgRNAs in Ciona. **a)** Mutagenesis efficacy rate of each sgRNA plotted against the same sgRNA’s Fusi/Doench algorithm score, obtained from CRISPOR. The Spearman’s rank correlation coefficient (rho) is 0.435 (p = 3.884e-05). **b)** sgRNAs grouped by sorted Fusi/Doench predicted scores (left: >60; right: <50). 18 of 23 sgRNAs of Fusi/Doench score over 60 showed a measured mutagenesis efficacy (MUT%) over 24%, classified as “good” (shaded green). In contrast, only 4 from the same set had a MUT% under 24%, classified as “bad”. “Good” and “bad” classifications were based on phenotype frequency in F0 (see text for details). Out of 25 sgRNAs with Fusi/Doench score under 50, 17 were “bad”, while only 8 were “good”. **c)** A Receiver Operating Characteristic (ROC) curve assesses the credibility of using Fusi/Doench score cutoff (from 0 to 100) to classify sgRNAs as either “good” (>24.5% efficacy) or “bad” (≤24.5% efficacy). Using Fusi/Doench cutoffs as such a classifier returns an AUC of 0.77 (black line), while an AUC of 0.5 (dashed red line) represents the performance of a classifier solely based on random chance. The optimal Fusi/Doench cutoff (above which a score is likely to indicate “good” sgRNAs) was found to be between 50 and 55. See **materials and methods** and **Supplementary Table 2** for details.

In a previous study, highly penetrant, tissue-specific loss-of-function phenotypes in F0 embryos were elicited using the Ebf.3 sgRNA (Stolfi *et al.* 2014), which in our current study had a measured mutagenesis efficacy of 37%. Further comparison of measured mutagenesis efficacy and mutant phenotype frequency for a series of *Ebf-*targeting sgRNAs revealed that an estimated sgRNA efficacy at least 25% correlated with over 70% reduction in the frequency of embryos expressing an *Islet* reporter construct (see below and **Figure 6**). We thus reasoned that a mutagenesis efficacy of ˜25% would be the minimum threshold of acceptable sgRNA activity for loss-of-function assays in F0. Within our dataset, among the sgRNAs that had a Fusi/Doench score >60, 18 of 23 (78%) had a mutagenesis efficacy rate over 24% (**Figure 3b**). In contrast, only 8 of 25 (32%) of sgRNAs with a Fusi/Doench score <50 had an efficacy over 24% (**Figure 3b**). Indeed, a receiver operating characteristic (ROC) analysis showed an area under the curve (AUC) of 0.77 when using Fusi/Doench score as a classifier of “good” (>24.5% efficacy) vs. “bad” (≤24.5%) sgRNAs (**Figure 3c**). The most accurate Fusi/Doench score cutoffs appeared to be between 50 and 55, when taking both specificity and sensitivity into account (**Supplementary Table 2, Supplementary Figure 2**). However, if the number of candidate sgRNAs is not limiting, it may be more desirable to use a Fusi/Doench score cutoff of 60 in order to avoid false positives (i.e., sgRNAs that are predicted as “good” when in reality they are “bad”). Thus, for *Ciona*, we recommend using CRISPOR to select sgRNAs with Fusi/Doench scores >60 and avoid those with Fusi/Doench <50. We believe this approach will significantly streamline the search for suitable U6 promoter-driven sgRNAs targeting one’s gene of interest.

### Multiplexed targeting with CRISPR/Cas9 generates large deletions in the Ciona genome

Large deletions of up to 23 kb of intervening DNA resulting from NHEJ between two CRISPR/Cas9-induced DSBs have been reported in *Ciona* (Abdul-Wajid *et al.* 2015). For functional analyses of protein-coding genes, such deletions would more likely produce null mutations than small deletions resulting from the action of lone sgRNAs. To test whether we could cause tissue-specific large deletions in F0 embryos, we targeted the forkhead/winged helix transcription-factor-encoding gene *Foxf* (**Figure 4a**), which contributes to cardiopharyngeal development in *Ciona* (Beh et al. 2007). We co-electroporated *Eef1a1>nls::Cas9::nls* with sgRNA vectors Foxf.4 and Foxf.2 (with induced mutagenesis rates of 39% and 18%, respectively). We extracted genomic DNA from electroporated embryos and PCR-amplified the sequence spanning both target sites. We obtained a specific ˜300 bp PCR product corresponding to the amplified region missing the ˜2.1 kbp of intervening sequence between the two target sites (**Figure 4b**). Cloning the deletion band and sequencing individual clones confirmed that the short PCR product corresponds to a deletion of most of the *Foxf* first exon and 5’ *cis-*regulatory sequences (Beh *et al.* 2007). We did not detect this deletion using genomic DNA extracted from embryos electroporated with either sgRNA alone. Similar deletion PCR products were observed, cloned, and sequenced for other genes including *Nk4, Fgfr, Mrf, Htr7-related, Bmp2/4,* and *Hand*, using pairs of highly mutagenic sgRNAs (**Supplementary Figure 3**). The largest deletion recorded was ˜3.6 kbp, with sgRNAs Nk4.2 (46% efficacy) and Nk4.3 (38% efficacy), entirely removing the sole intron of *Nk4* and small portions of the flanking exons. The sgRNAs targeting *Mrf* were shown to inhibit its function and subsequent siphon muscle development (Tolkin and Christiaen 2016).

**Figure 4.**
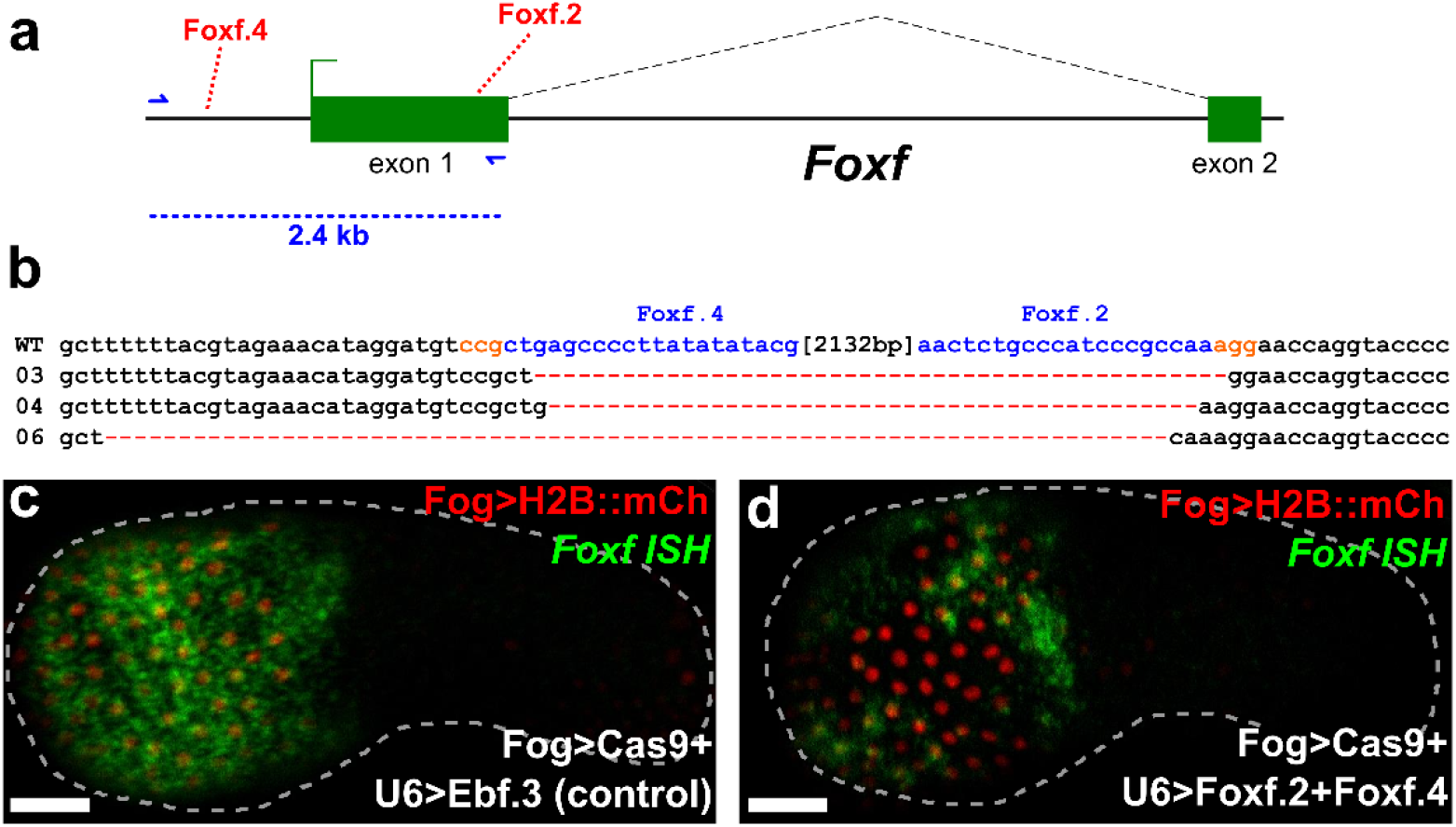
Combinatorial targeting of Foxf results in large deletions. **a)** Diagram of *Foxf* locus, showing positions targeted by *Foxf.4* and *Foxf.2* sgRNAs. *Foxf.4* targets a non-coding, *cis-*regulatory sequence 881 base pairs (bp) upstream of the transcription start site of *Foxf. Foxf.2* targets a coding sequence in exon 1 of *Foxf.* The distance between the target sites is 2132 bp, and encompasses most of exon 1, the core promoter, and *cis-*regulatory modules that drive *Foxf* expression in the head epidermis and trunk ventral cells (TVCs) (Beh *et al.* 2007). Blue arrows indicate primers used to amplify the region between the target sites. In wild-type embryos, the resulting PCR product is ˜2.4 kilobase pairs (kbp). **b)** Alignment of cloned PCR products amplified using the primers indicated in (a), from wild-type (wt) embryos, and from embryos electroporated with 25µg *Eef1a1>nls::Cas9::nls* and 50 µg each of *U6>Foxf.2* and *U6>Foxf.4.* Colonies 03, 04, and 06 shown containing large deletions between the approximate sites targeted by the two sgRNAs, indicating non-homologous end-joining (NHEJ) repair from two separate double stranded break events as a result of combinatorial action of *Foxf.2* and *Foxf.4* sgRNAs. **c)** *In situ* hybridization for *Foxf* (green) showing strong expression throughout the head epidermis in embryos electroporated with 10 µg *Fog>H2B::mCherry* (red), 50 µg *Fog>nls::Cas9::nls* and 45 µg of *U6>Ebf.3. Foxf* expression is essentially wild-type, as Ebf function is not required for activation of *Foxf* in the epidermis. **d)** *In situ* hybridization for *Foxf* (green) showing patchy expression in the head epidermis of embryos electroporated with 10 µg *Fog>H2B::mCherry* (red), 50 µg *Fog>nls::Cas9::nls* and 45 µg each of *U6>Foxf.2* and *U6>Foxf.4*. Loss of *in situ* signal in some transfected head epidermis cells indicates loss of *Foxf* activation, presumably through deletion of all or part of the upstream *cis*-regulatory sequences by CRISPR/Cas9. Scale bars = 25 µm.

*Foxf* is expressed in TVCs and head epidermis (**Figure 4c**), the latter of which is derived exclusively from the animal pole (Nishida 1987; Imai *et al.* 2004; Pasini *et al.* 2006; Beh *et al.* 2007). Because the ˜2.1 kbp deletion introduced in the *Foxf* locus excised the epidermal enhancer and basal promoter (Beh *et al.* 2007), we sought to examine the effects of these large deletions on *Foxf* transcription. We used the *cis*-regulatory sequences from *Zfpm* (also known as *Friend of GATA*, or *Fog*, and referred to as such from here onwards) to drive Cas9 expression in early animal pole blastomeres (Rothbächer *et al.* 2007). We electroporated *Fog>nls::Cas9::nls* together with Foxf.2 and Foxf.4 sgRNA vectors and *Fog>H2B::mCherry*, and raised embryos at 18°C for 9.5 hpf (early tailbud, embryonic stage 20). We performed whole mount mRNA *in situ* hybridization to monitor *Foxf* expression, expecting it to be silenced in some epidermal cells by tissue-specific CRISPR/Cas9-induced deletions of the *Foxf cis-* regulatory sequences on both homologous chromosomes in each cell. Indeed, we observed patches of transfected head epidermal cells (marked by H2B::mCherry) in which *Foxf* expression was reduced or eliminated (**Figure 4d**). This was in contrast to the uniform, high levels of *Foxf* expression observed in “control” embryos electroporated with Ebf.3 sgRNA (*Ebf* is unlikely to be involved in *Foxf* regulation in the epidermis where it is not expressed, **Figure 4c**). Taken together, these results indicate that, by co-electroporating two or more highly active sgRNAs targeting neighboring sequences, one can frequently generate large deletions in the *Ciona* genome in a tissue-specific manner.

### Rapid generation of sgRNA expression cassettes ready for embryo transfection

CRISPR/Cas9 is an efficient and attractive system for targeted mutagenesis in *Ciona*, but cloning individual sgRNA vectors is a labor-intensive, rate-limiting step. To further expedite CRISPR/Cas9 experiments, we adapted a one-step overlap PCR (OSO-PCR) protocol to generate U6 promoter>sgRNA expression “cassettes” for direct electroporation without purification (**Figure 5, Supplementary Figure 4**, see **Materials and Methods** and **Supplementary Protocol** for details). We tested the efficacy of sgRNAs expressed from these unpurified PCR products, by generating such expression cassettes for the validated Ebf.3 sgRNA. We electroporated *Eef1a1>nls::Cas9::nls* and 25 µl (corresponding to ˜2.5 µg, see **Materials and Methods** and **Supplementary Figure 5**) of unpurified, *U6>Ebf.3* sgRNA OSO-PCR or *U6>Ebf.3* sgRNA traditional PCR products (total electroporation volume: 700 µl). Next-generation sequencing of the targeted Ebf.3 site revealed mutagenesis rates similar to those obtained with 75 µg of *U6>Ebf.3* sgRNA plasmid (**Supplementary Table 1**). This was surprising given the much lower total amount of DNA electroporated from the PCR reaction relative to the plasmid prep (2.5 µg vs. 75 µg). This discrepancy could indicate a higher efficiency of transcription of linear vs. circular transfected DNA, though more thorough analyses are warranted to investigate the behavior of linear DNA in *Ciona* embryos.

**Figure 5.**
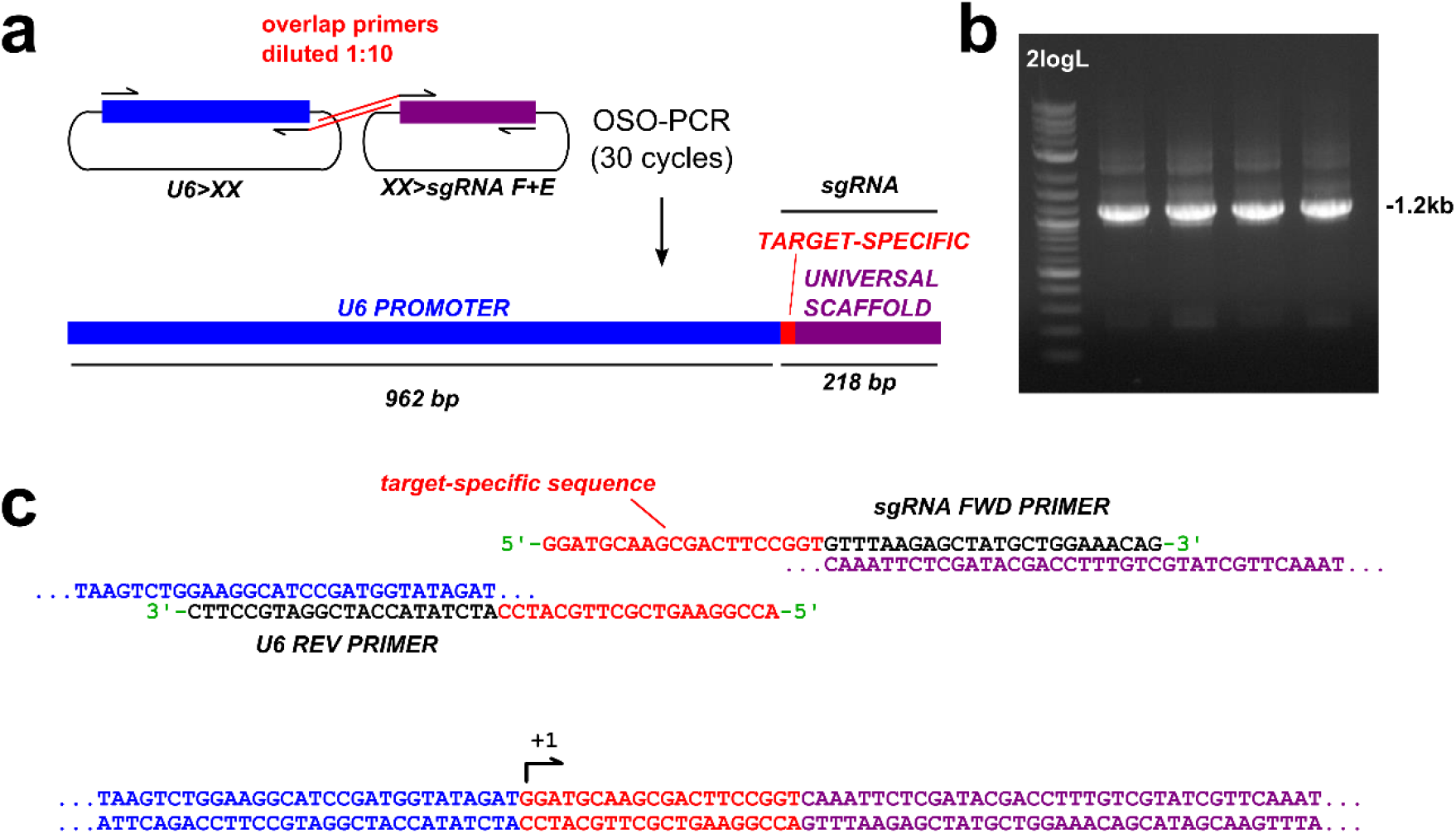
One-step Overlap Polymerase Chain Reaction (OSO-PCR) for the high-throughput construction of sgRNA expression cassette libraries. **a)** Diagram of OSO-PCR for amplification of *U6>sgRNA* expression cassettes in which the target-specific sequence of each (red) is encoded in complementary overhangs attached to universal primers. 1:10 dilution of these primers ensures that the overlap product, the entire U6>sgRNA cassette, is preferentially amplified (see methods for details). **b)** Agarose gel electrophoresis showing products of four different *U6>sgRNA* OSO-PCRs. The desired product is ˜1.2 kilobase pairs (kbp) long. 2logL = NEB 2-Log DNA ladder. **c)** Detailed diagram of how the overlap primers form a target-specific bridge that fuses universal U6 promoter and sgRNA scaffold sequences.

To assess whether unpurified sgRNA PCR cassettes could be used in CRISPR/Cas9-mediated loss-of-function experiments in F0 embryos, we assayed the expression of an *Islet* reporter transgene in MN2 motor neurons (Ryan *et al.* 2016), which depends upon Ebf function (Stolfi *et al.* 2014). Indeed, *Islet>GFP* expression was downregulated in embryos electroporated with *Sox1/2/3>nls::Cas9::nls* and 25 µl of unpurified *U6>Ebf.3* traditional PCR or 25µg *U6>Ebf.3* plasmid, but not with 25 µl (˜2.5 µg) of unpurified *U6>Negative Control* sgRNA PCR cassette (**Figure 6a-d**). Taken together, these results indicate that unpurified PCR products can be used in lieu of plasmids to express sgRNAs for tissue-specific CRISPR/Cas9-mediated mutagenesis in *Ciona* embryos.

**Figure 6.**
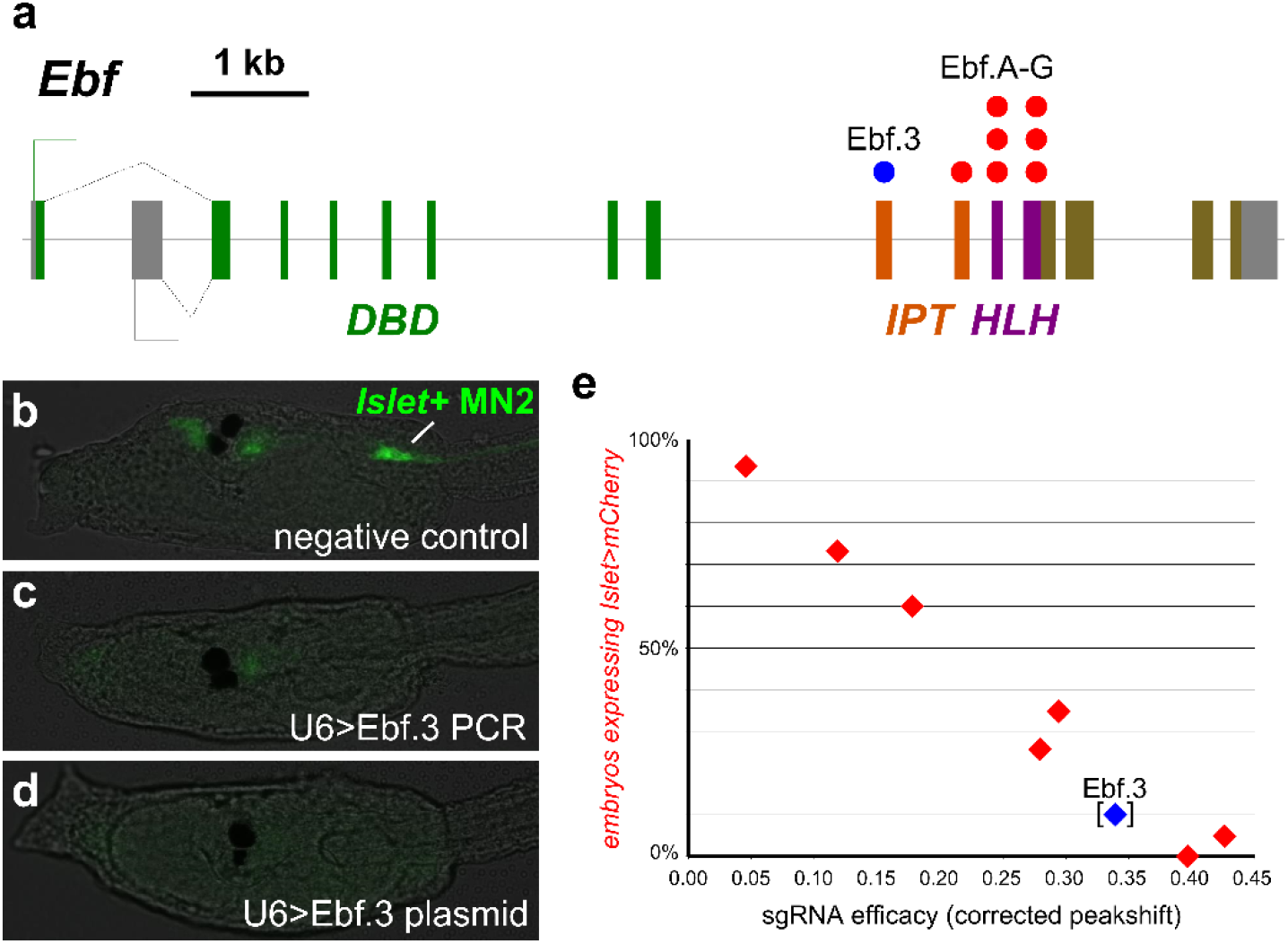
Linear relationship between sgRNA efficacy and mutant phenotype frequency in F0. **a)** Diagram of *Ebf* locus, showing exons coding for DNA-binding (DBD), IPT, and helix-loop-helix (HLH) domains. Colored dots indicate exons targeted by the *Ebf-*targeting sgRNAs used to validate the OSO-PCR method for genetic loss-of-function. **b)** Larvae co-electroporated with *Sox1/2/3>nls::Cas9::nls, Islet>eGFP*, and 25 µl (˜2.5 µg) unpurified *U6>NegativeControl* PCR or **c)** 25 µl (˜2.5 µg) unpurified *U6>Ebf.3* PCR, or **d)** 25 µg *U6>Ebf.3* plasmid. *Islet>eGFP* reporter plasmid is normally expressed in MN2 motor neurons (“Islet+ MN2”, green), which is dependent on Ebf function. *Islet>eGFP* was expressed in MN2s in 75 of 100 negative control embryos. In embryos electroporated with unpurified *U6>Ebf.3* PCR products or *U6>Ebf.3* plasmid, only 16 of 100 and 17 of 100 embryos, respectively, had *Islet>eGFP* expression in MN2s. This indicates similar loss of Ebf function *in vivo* by either unpurified PCR or purified plasmid sgRNA delivery method. **c)** Plot comparing mutagenesis efficacies of the OSO-PCR-generated sgRNAs indicated in panel (a) (measured by Sanger sequencing, see text for details) and the ability to cause the *Ebf* loss-of-function phenotype (loss of *Islet>mCherry* reporter expression in MN2s in *Sox1/2/3>H2B::eGFP+* embryos). The nearly perfect inverse correlation between sgRNA mutagenesis efficacy and *Islet>mCherry* expression indicates a linear relationship between sgRNA activity and mutant phenotype frequency in electroporated embryos. Ebf.3 sgRNA data point is bracketed, because its mutagenesis efficacy was not measured by Sanger sequencing but comes from the NGS data collected in this study.

### Correlation between sgRNA efficacy and mutant phenotype penetrance

Since we measured a wide range of mutagenesis efficacies across our library of 83 sgRNAs, we wanted to assess how this variability correlates with mutant phenotype penetrance. One possibility is that sgRNA efficacies and frequencies of mutant phenotypes in F0 are linearly correlated. Alternatively, mutant phenotypes might only be observed at distinct thresholds of sgRNA activity.

To test this, we designed 7 additional sgRNAs (Ebf.A through Ebf.G) targeting the IPT and HLH domain-coding exons of *Ebf* (**Figure 6a,e, Supplementary Table 3**). We generated these sgRNA expression cassettes by OSO-PCR, and co-electroporated 25 µl of each reaction with either 25 µg *Eef1a1>nls::Cas9::nls* (for mutagenesis efficacy sequencing) or 40 µg *Sox1/2/3>nls::Cas9::nls +* 15 µg *Sox1/2/3>H2B::eGFP* + 40 µg *Islet>mCherry* (for phenotype assay). sgRNA efficacies were measured by direct Sanger sequencing of the target region PCR-amplified from larvae electroporated with the former combination (See **Materials and methods** for details), and *Islet>mCherry* expression in H2B::eGFP+ motor neurons was scored in larvae electroporated with the latter mix.

Quantification of indel-shifted electrophoresis chromatogram peaks (“Peakshift” assay) revealed sgRNA mutagenesis efficacies ranging from 5% to 43% (**Figure 6e, Supplementary Table 3**). Of note, the sgRNA with the lowest efficacy (Ebf.E) was clearly hampered by a naturally occurring SNP that eliminates the PAM (NGG to NGA) in a majority of haplotypes. In parallel, the proportion of transfected embryos *Islet>mCherry* expression was observed to vary between 0% and 94%. When this was plotted against the measured mutagenesis efficacies, we observed a nearly perfect linear correlation between the mutant phenotype (loss of *Islet>mCherry* expression) and *Ebf* mutagenesis (**Figure 6e, Supplementary Table 3**). Taken together, these results suggest that highly efficient sgRNAs can be generated from OSO-PCR cassettes and validated for loss-of-function studies by Sanger sequencing. The Sanger sequencing-based peakshift assay is highly reproducible and approximates the efficacy rates estimated by NGS (**Supplementary Table 6**). As such, we recommend the peakshift assay as a cheap, fast alternative for testing sgRNA activity in *Ciona* embryos.

### Genome-wide design and scoring of sgRNAs for Ciona

While *Ciona* researchers will find the Fusi/Doench scoring system and the OSO-PCR method useful for sgRNA expression cassette design and assembly, we hoped to further empower the community by pre-emptively designing all possible sgRNAs with a unique target site within 200 bp of all *Ciona* exonic sequences. We computationally identified 3,596,551 such sgRNAs. We have compiled all sgRNA sequences and their corresponding specificity and efficiency scores (including Fusi/Doench). They are available as a UCSC Genome Browser track for the Ci2 genome assembly (http://genome.ucsc.edu/), with links to CRISPOR for off-target prediction and automated OSO-PCR primer design. This track is freely available for download to use on other browsers, like those of specific interest to the tunicate community such as ANISEED (Brozovic *et al.* 2015) or GHOST (Satou *et al.* 2005).

## Discussion

We have built a library of 83 plasmid vectors for the *in vivo* expression of sgRNAs targeting 23 genes expressed in the cardiopharyngeal mesoderm and surrounding tissues, mostly hypothesized to be involved in regulating the specification of heart and/or pharyngeal muscles in *Ciona*, even though many have complex expression patterns and probably pleiotropic functions. We have also established reliable protocols for the validation of sgRNA efficacy in electroporated *Ciona* embryos by either next-generation sequencing or Sanger sequencing. This has allowed us to estimate the activity of all these sgRNAs, which are ready to be used for ongoing and future functional studies. We are aware that the lack of biological replicates for the NGS-based measurements and other technical limitations of either assay may affect the accuracy of some of our mutagenesis efficacy estimates. However, our observations suggest that our results were not greatly affected by electroporation variability (**Supplementary Figure 1; Supplementary Table 6**), and that our data are accurate enough to identify sgRNAs of high or low activity in *Ciona*.

By analyzing correlations between target sequence nucleotide composition and sgRNA mutagenesis efficacy, we identified sgRNA sequence features that may contribute to Cas9:sgRNA activity. Some of these sequence features have been identified in previous CRISPR/Cas9-mediated mutagenesis screens performed in other metazoan model organisms, suggesting that these are determined by the intrinsic properties of sgRNAs and/or Cas9 (Doench *et al.* 2014; Gagnon *et al.* 2014; Ren *et al.* 2014; Chari *et al.* 2015; MORENO-Mateos *et al.*2015). For instance, sgRNA efficacy is correlated with increased guanine content in the PAM-proximal nucleotides of the sgRNA, postulated to be due to increased sgRNA stability by G-quadruplex formation (MORENO-Mateos *et al.* 2015). This would explain the specific enrichment for guanine but not cytosine, even if both could in theory augment sgRNA folding or binding to target DNA. We also encountered a depletion of thymine and cytosine in the PAM-proximal nucleotides of the protospacers for highly active sgRNAs. The strong negative correlation between sgRNA efficacy and thymine content of the protospacer is easily attributed to our use of the PolIII-dependent U6 promoter to express our sgRNAs. It has been shown that termination of transcription by PolIII can be promoted by degenerate poly-dT tracts (Nielsen *et al.* 2013). A high number of non-consecutive thymines clustered in the protospacer could thus result in lower sgRNA expression level due to premature termination of sgRNA transcription (Stolfi *et al.* 2014; Wu *et al.* 2014). Similarly, adenines are thought to contribute to the instability of sgRNAs (MORENO-Mateos *et al.* 2015), suggesting that CRISPR/Cas9 mutagenesis efficacies might be primarily determined by sgRNA transcription and degradation rates, which will vary depending on the species studied and the mode of sgRNA delivery (e.g. *in vitro* vs. *in vivo* synthesis).

We demonstrate that among published sgRNA prediction algorithms, Fusi/Doench (Fusi *et al.* 2015; Doench *et al.* 2016) functions well as a classifier for “good” (>24.5% mutagenesis efficacy) and “bad” (<24.5%) sgRNAs. Despite these general trends, several sgRNAs defied this algorithm-based prediction. This suggests that there are multiple, possibly additive or synergistic factors that determine the mutagenesis efficacy, only one of which is primary sequence composition of the sgRNA or target. Other factors that can influence Cas9 binding include chromatin accessibility and nucleosome occupancy (Wu *et al.* 2014; Hinz *et al.* 2015; Horlbeck *et al.* 2016; Isaac *et al.* 2016). What other additional factors contribute to variability of *in vivo* sgRNA mutagenesis efficacies will be an important topic of study as CRISPR/Cas9-based approaches are expanded to address additional questions in basic research as well as for therapeutic purposes.

While an optimized predictive algorithm for *Ciona*-specific sgRNA design remains a desirable goal, our current approach should help other researchers to identify, with greater confidence,which sgRNAs are likely to confer enough mutagenic activity for functional studies in F0. We have shown, using a series of *Ebf*-targeting sgRNAs and an Ebf loss-of-function readout assay (*Islet* reporter expression) that a linear correlation exists between sgRNA mutagenesis efficacy and the probability of somatic gene knockout in F0. In this case, we measured the frequency of a binary loss-of-function assay (*Islet* ON or OFF) in a large population of embryos. Therefore it is expected that the probability a homozygous *Ebf* knockout (resulting in *Islet* OFF phenotype) should be correlated to the observed frequency of *Ebf* mutations in somatic cell populations. While Ebf function can be disrupted by small changes to its crucial IPT/HLH domains, other genes may prove harder to disrupt with the short indels generated by CRISPR/Cas9. However, we show that the combinatorial action of two or more sgRNAs can result in high frequency of large deletions spanning many kbp, which should help generate loss-of-function alleles for such recalcitrant genes.

Despite legitimate concerns about potential off-target effects for functional studies, we were not able to detect CRISPR/Cas9-mediated mutagenesis at two potential off-target sites for the sgRNAs Ebf.3 and Fgf4/6.1. For the remainder of the sgRNAs, we purposefully selected those with no strongly predicted off-targets. This was possible in *Ciona* due to two factors. First, the *Ciona* genome is significantly smaller than the human genome and most metazoans, resulting in a lower number of similar protospacer sequences. Second, the GC content of the *Ciona* genome is only 35% as compared to 65% in humans, which should result in a lower overall frequency of canonical PAMs (NGG). Based on these considerations, we predict off-target effects to be less pervasive in *Ciona* than in other model organisms with more complex, GC-rich genomes.

Even with improved prediction of sgRNA efficacy and specificity, there is still a need to test several sgRNAs to identify the optimal one targeting a gene of interest. To this end, we have developed a cloning-free OSO-PCR method for the rapid assembly of sgRNA expression cassettes. We have shown that these unpurified PCR cassettes can be directly electroporated into *Ciona* embryos and screened by either Sanger sequencing of target sequences or mutant phenotype frequency in F0. The automated design of primers for Ciona-specific sgRNA cassette OSO-PCR has been implemented in the latest version of CRISPOR (http://crispor.tefor.net/).

Finally, we have pre-emptively designed all possible sgRNAs targeting exonic sequences in the compact *Ciona* genome and calculated their specificity and efficiency by various predictive algorithms. This track is available online on the UCSC genome browser, but is also freely available for download and use with other genome browsers. This allows researchers to locally browse for sgRNAs with predicted high activity targeting their loci of interest. Integration with the CRISPOR website further allows SNP and off-target prediction, and pre-designed OSO-PCR oligonucleotide primers for rapid, efficient synthesis and transfection. We expect these resources to facilitate the scaling of CRISPR/Cas9-mediated targeted mutagenesis and enable genome-wide screens for gene function in *Ciona*.

## Materials and Methods

### Target sequence selection and sgRNA design

23 genes from *Ciona robusta* (formerly *Ciona intestinalis* type A)(Hoshino and Tokioka 1967; Brunetti *et al.* 2015) hypothesized to be important for cardiopharyngeal mesoderm development were shortlisted (**Table 1**) and one to four sgRNAs targeting non-overlapping sequences per gene were designed, for a total of 83 sgRNA vectors (**Supplementary Table 3**). Two sgRNAs were designed to target the neurogenic bHLH factor Neurogenin, a gene that is not expressed in the cardiopharyngeal mesoderm and is not thought to be involved in cardiopharyngeal development. Target sequences were selected by searching for N19 + NGG (protospacer + PAM) motifs and screened for polymorphisms and off-target matches using the GHOST genome browser and BLAST portal (Satou *et al.* 2005; Satou *et al.* 2008). Potential off-targets were also identified using the CRISPRdirect platform (Naito *et al.* 2015). sgRNA expression plasmids were designed for each of these protospacers and constructed using the *U6>sgRNA(F+E)* vector as previously described (Stolfi *et al.* 2014), as well as a “Negative Control” protospacer that does not match any sequence in the *C. robusta* genome (5’-GCTTTGCTACGATCTACATT-3’). Stretches of more than four thymine bases (T) were avoided due to potential premature transcription termination. Candidate sgRNAs with a partial PAM-proximal match of 13 bp or more to multiple loci were also discarded due to off-target concerns. All sgRNAs were designed to target protein-coding, splice-donor, or splice-acceptor sites, unless specifically noted. We preferred more 5’ target sites, as this provides a greater probability of generating loss-of-function alleles.

### Electroporation of Ciona embryos

DNA electroporation was performed on pooled, dechorionated zygotes (1-cell stage embryos) from *C. robusta* adults collected from San Diego, CA (M-REP) as previously described (Christiaen *et al.* 2009). All sgRNA plasmid maxipreps were individually electroporated at a final concentration of 107 ng/µl (75 µg in 700 µl) concentration together with *Eef1a1>nls::Cas9::nls* plasmid (Stolfi *et al.* 2014) at 35.7 ng/µl (25 µg in 700 µl)concentration. For testing *U6>Ebf.3* PCR or OSO-PCR, 25 µl was used instead of sgRNA plasmid. Embryos were then rinsed once in artificial sea water, to remove excess DNA and electroporation buffer, and grown at 18°C for 16 hours post-fertilization.

### Embryo lysis

After 16 hpf, each pool of embryos targeted with a single sgRNA + Cas9 combination was washed in one sea water exchange before lysis, to remove excess plasmid DNA, and transferred to a 1.7 ml microcentrifuge tube. Excess sea water was then removed and embryos were lysed in 50 µl of lysis mixture prepared by mixing 500 µL of DirectPCR Cell Lysis Reagent (Viagen Biotech Inc., Los Angeles, CA, Cat # 301-C) with 1 µl of Proteinase K (20 mg/ml, Life Technologies, Carlsbad, CA). The embryos were thoroughly mixed in lysis mixture and incubated at 68°C for 15 minutes, followed by 95°C for 10 minutes.

### PCR amplification of targeted sequences

Targeted sequences were individually PCR-amplified directly from lysate from embryos targeted with the respective sgRNA, and from “negative control” lysate (from embryos electroporated with *Eef1a1>nls::Cas9::nls* and *U6>Negative Control* sgRNA vector). Primers (**Supplementary Table 4**) were designed to flank target sites as to obtain PCR products in the size range of 108-290bp with an exception of the sequence targeted by Ebf.3 (“Ebf.774” in Stolfi et al. 2014) and Ebf.4 sgRNAs, for which the designed primers resulted in a product size of 350 bp. Potential off-target sites predicted for sgRNAs Ebf.3 (CTCGCAACGGGGACAACAGGGGG, genome position KhC8:2,068,844-2,068,866) and Fgf4/6.1 (TATTTTAATTCTGTACCTGTGGG, genome position KhC9:6,318,421-6,318,443) were amplified to test for off-target CRISPR/Cas9 activity with the primers: 5’-CCAGCACTTCAGAGCAATCA-3’ and 5’-TGACGTCACACTCACCGTTT-3’ (Ebf.3), and 5’-AACGATTGTCCATACGAGGA-3’ and 5’-ACTTCCCAACAGCAAACTGG-3’ (Fgf/6.1).

For each targeted sequence, 12.5 µL PCR reactions were set up with final concentrations of 600 nM each primer, 300 µM dNTPs, 1 mM MgSO4, 2X buffer, and 0.05 U/µl Platinum Pfx DNA polymerase (Life Technologies), and subjected to the following PCR program: an initial cycle of 10 minutes at 95°C, followed by 30 cycles of 30 seconds at 94°C, 30s at 60°C and 30s at 68°C, and a final cycle of 3 minutes at 68°C. PCR reactions were quickly checked on an agarose gel for the presence/absence of amplicon. Those that resulted in a single band were not initially purified. For those reactions with more than one band, the correct amplicon (selected based on expected size) was gel purified using a Nucleospin Gel Clean-up Kit (Macherey-Nagel, Düren, Germany). Purified and unpurified PCR products were then pooled for subsequent processing. The majority of PCR products amplified from larvae treated with Cas9 + gene-targeting sgRNA were pooled in Pool 1. All products from larvae treated with Cas9 + “negative control” sgRNA were pooled in Pool 2. For those sequences targeted by distinct sgRNAs but amplified using the same set of flanking primers, their PCR products were split into separate pools, as to allow for separate efficacy estimates.

### Sequencing library preparation

The PCR product pools were electrophoresed on ethidium bromide-stained, 1% agarose gel in 0.5X Tris-Acetate-EDTA (TAE) buffer and a band of ˜150-300 bp was excised (Nucleospin gel and PCR cleanup kit, Macherey-Nagel). 102-235 ng of each pool was used as a starting material to prepare sequencing libraries (protocol adapted from http://wasp.einstein.yu.edu/index.php/Protocol:directional_WholeTranscript_seq). Ends were repaired using T4 DNA polymerase (New England Biolabs, Ipswich, MA) and T4 Polynucleotide Kinase (New England Biolabs), and then A-tailed using Klenow fragment (3'→5' exo-) (New England Biolabs) and dATP (Sigma-Aldrich, St. Louis, MO). Each pool was then ligated to distinct barcoded adapters. (NEXTflex DNA Barcodes - BioO Scientific Cat# 514101). The six barcodes used in this study were: CGATGT, TGACCA, ACAGTG, GCCAAT, CAGATC and CTTGTA. At this step, the adapter-ligated DNA fragments were purified twice using Ampure XP beads (Beckman Coulter, Brea, CA). The final amplification, using primers included with NEXTflex adapters, was done using the PCR program: 2 minutes at 98°C followed by 8 cycles of 30 seconds at 98°C; 30 seconds at 60°C; 15 seconds of 72°C, followed by 10 minutes at 72°C. Ampure XP bead-based selection was performed twice, and the libraries were quantified using qPCR. The libraries were then mixed in equimolar ratio to get a final DNA sequencing library concentration of 4 nM. The multiplexed library was sequenced by Illumina MiSeq V2 platform (Illumina, San Diego, CA) using 2x250 paired end configuration.

### Next generation sequencing data analysis

FastQ files obtained from sequencing were de-multiplexed and subjected to quality control analysis. FastQ reads were mapped to the 2008 KyotoHoya genome assembly (Satou *et al.* 2008) by local alignment using Bowtie2.2 (Langmead and Salzberg 2012). Single end reads were also mapped to a reduced genome assembly consisting of only those scaffolds containing the targeted genes. This allowed for a much faster and accurate alignment using Bowtie2.2. The SAM file generated was converted into a BAM file using *samtools* (Li *et al.* 2009). The BAM file was sorted and indexed to visualize reads on Integrative Genomics Viewer (IGV) (Robinson *et al.* 2011). Most mutagenesis rates were obtained by counting indels in IGV. For some targets with partially overlapping aplicon sequences, custom Python scripts were written to parse the BAM file to get estimated rate of mutagenesis. Since a successful CRISPR/Cas9-mediated deletion or insertion should eliminate or disrupt all or part of the protospacer + PAM sequence (jointly termed the “pineapple”), we simply looked for mapped reads in which the pineapple was not fully present. When appropriate, the rate of naturally occurring indels around each target, as detected in reads from “negative control” embryos, was subtracted from the raw efficacy rates. Custom python scripts used are available upon request. Matplotlib (http://matplotlib.org) was used for plotting, Numpy (http://numpy.org) and Pandas (http://pandas.pydata.org) were used for data mining. All predictive algorithm scores were generated using CRISPOR (http://crispor.tefor.net/)(Haeussler *et al.* 2016).

### Nucleotide Enrichment Analysis

We used log-odds score as a measure to estimate how enriched each nucleotide was at a given position in the 43bp region of interest (excluding position 1 of the protospacer, and the ‘GG’ of the PAM). Log-odds score was defined as the base-2 logarithm of the ratio of probability of observing nucleotide ‘N’ at position ‘x,’ and the background probability of observing nucleotide ‘N’ by random chance, given its frequency in our sample space (all sgRNA targets, n=83). A positive or negative log-odds score reflects enrichment or depletion, respectively, of each nucleotide at a given position.

Mathematically,

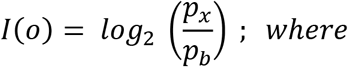

*p_x_* = probability of finding nucleotide ‘N’ at position ‘x’
*p_b_* = background probability of finding nucleotide ‘N’ at position ‘x’ by chance

### Receiver Operating Characteristic (ROC) curve

For a binary classifier (“good” vs. “bad” sgRNA), there are four possible outcomes: True Positive (TP), False Positive (FP), True Negative (TN), and False Negative (FN). If the predicted classification of the sgRNA by our model was “good” or “bad”, and it was supported by our experimental data (i.e. the empirical mutagenesis rate was above or below 24.5% respectively), it was marked as a “True Positive” or a “True Negative,” respectively. If the experimental data failed to support it, it was marked as a “False Positive” or a “False Negative,” respectively. The true positive rate (TPR), or “Sensitivity”, was defined as the proportion of empirically good sgRNAs that were correctly predicted as good [TPR = TP / (TP + FN)]. Similarly, the true negative rate (TNR), or “Specificity”, was defined as the proportion of empirically bad gRNAs that were correctly predicted as bad [TNR = TN / (TN + FP)]. The false positive rate (FPR) is 1 - Specificity. In figure **3c**, we plotted the true positive rates against the false positive rates, all obtained by applying different Fusi/Doench score thresholds (ranging from 0 to 100) to the predictions generated by our model (see **Supplementary Table 2**).

### Combinatorial sgRNA electroporation to induce large deletions

Embryos were electroporated with 25 µg *Eef1a1>nls::Cas9::nls* and two vectors from the set of validated sgRNA plasmids for each targeted gene (50 µg per sgRNA vector). Embryos were grown for 12 hpf at 18°C, pooled, and genomic DNA extracted from them using QIAamp DNA mini kit (Qiagen). Deletion bands were amplified in PCR reactions using Pfx platinum enzyme as described above (see “**PCR amplification of targeted sequences**”) and a program in which the extension time was minimized to 15 seconds only, in order to suppress the longer wild-type amplicon and promote the replication of the smaller deletion band. Primers used were immediately flanking the sequences targeted by each pair of sgRNAs (**Supplementary Table 4**). Products were purified from agarose gels, A-overhung and TOPO-cloned. Colonies were picked, cultured, prepped and sequence

### Synthesis and electroporation of unpurified sgRNA PCR expression cassettes

*U6>Ebf.3* and *U6>Negative Control* sgRNA expression cassettes were amplified from their respective plasmids using the primers U6 forward (5’-TGGCGGGTGTATTAAACCAC-3’) and sgRNA reverse (5’-GGATTTCCTTACGCGAAATACG-3’) in reactions of final concentrations of 600 nM each primer, 300 µM dNTPs, 1 mM MgSO4, 2X buffer, and 0.05 U/µl Platinum Pfx DNA polymerase (Life Technologies), and subjected to the following PCR program: an initial cycle of 3 minutes at 95°C, followed by 30 cycles of 30 seconds at 94°C, 30s at 55°C and 2 minutes at 68°C, and a final cycle of 5 minutes at 68°C. *U6>sgRNA(F+E)::eGFP* (Stolfi *et al.* 2014) was amplified as above but using Seq forward primer (5’-AGGGTTATTGTCTCATGAGCG-3’) instead. For phenotyping Ebf-dependent expression of *Islet* reporter in MN2 motor neurons (Stolfi *et al.* 2014), embryos were co-electroporated with 35-40 µg of *Sox1/2/3> nls::Cas9::nls,* 5-15 µg of *Sox1/2/3>H2B::mCherry/eGFP,* 30-40 µg of *Isl>eGFP/mCherry*, and either 25 µg of *U6>Ebf.3* plasmid or 25 µl (˜2.5 µg) of unpurified PCR product.

### sgRNA expression cassette assembly by One-step Overlap PCR (OSO-PCR)

sgRNA PCR cassettes were constructed using an adapted One-step Overlap PCR (OSO-PCR) protocol (Urban *et al.* 1997). Basically, a desired protospacer sequence is appended 5’ to a forward primer (5’-GTTTAAGAGCTATGCTGGAAACAG-3’) and its reverse complement is appended 5’ to a reverse primer (5’-ATCTATACCATCGGATGCCTTC-3’). These primers are then added to a PCR reaction at limiting amounts, together with U6 forward (5’-TGGCGGGTGTATTAAACCAC-3’) and sgRNA reverse (5’-GGATTTCCTTACGCGAAATACG-3’) primers and separate template plasmids containing the U6 promoter (*U6>XX*) and the sgRNA scaffold (*XX>sgRNA F+E*). Plasmids are available from Addgene (https://www.addgene.org/Lionel_Christiaen/). The complementarity between the 5’ ends of the inner primers bridges initially separate U6 and sgRNA scaffold sequences into a single amplicon, and because they are quickly depleted, the entire cassette is preferentially amplified in later cycles by the outer primers (see **Figure 5** and **Supplementary Protocol** for details). Final, unpurified reactions should contain PCR amplicon at ˜100 ng/µl, as measured by image analysis after gel electrophoresis (**Supplementary Figure 5**).

### Measuring mutagenesis efficacies of sgRNAs by Sanger sequencing

OSO-PCR cassettes were designed and constructed for the expression of 7 sgRNA targeting exons 10-12 that code for part of the IPT and HLH domains of Ebf (**Figure 6a, Supplementary Table 3**). Embryos were electroporated with 25 µl of unpurified OSO-PCR product and 25 µg *Eef1a1>nls::Cas9::nls*, and grown to hatching. For the replication experiment (**Supplementary Table 6**) 75 µg of sgRNA vectors (Gata4/5/6.2, Gata4/5/6.3, Gata4/5/6.4, Neurog.1, Neurog.2) were each co-electroporated with 25 µg *Eef1a1>nls::Cas9::nls.* This was repeated over two separate batches of embryos.

All larvae were allowed to hatch and genomic DNA was extracted using the QiaAmp DNA mini kit (Qiagen). The *Ebf* target region was PCR-amplified using the following primers: (Forward primer: 5’-CTCCACATGCCTCAACTTTG-3’, Reverse primer: 5’-TGTTCCGCCAAATTGTGAAG-3’). For *Gata4/5/6* target sequences, the primers used were the corresponding ones from the NGS experiment, while a novel pair of primers was used to amplify the *Neurog* locus (Forward primer: 5’-AAGTACGGAGAGCAGAATACC-3’, Reverse primer: 5’-CTTCTAGTGCGTCATTAAGACC-3’). PCR was performed in 35 cycles using Platinum Pfx DNA polymerase. The resulting PCR products were gel-or column-purified, and sequenced using either a flanking PCR primer or an internal sequencing primer (Ebf internal: 5’-AATTGGCTGACAGGTTGGAG-3’, Neurog internal: 5’-GCTCTTGCTACAAAATGTTGG-3’).

The resulting .ab1 sequencing files were then analyzed by ab1 Peak Reporter webtool (Roy AND Schreiber 2014) (https://apps.thermofisher.com/ab1peakreporter/)(**Supplementary Table 5**). To quantify the peak “shifts” resulting from CRISPR/Cas9-induced short indels (**Supplementary Figure 6**), we calculated the sum of “maximum signal 7 scan filtered ratio” (MaxSig7Scan Sum) values of minor peaks at each position in a 30 bp window starting from the third bp in the target sequence from the PAM (the most likely Cas9 cut site). The mean MaxSig7Scan Sum was calculated across all 30 bp of this window, resulting in a “raw peakshift score”. The same was repeated for the 30 bp window immediately 5’ to the cut site in the sequencing read, for a peakshift “baseline” estimate. This baseline was subtracted from the raw peakshift score to give the “corrected peakshift score”, a relative quantification of indel frequency, and therefore of the mutagenesis efficacy of the sgRNA used.

### Whole-exome sgRNA predictions in *Ciona*

We used all transcribed regions in the ENSEMBL 65 transcript models (Aken *et al.* 2016), extended them by 200 bp on each side, searched for all possible NGG-20bp sgRNA targets in these sequences and ran them through the command-line version of CRISPOR (Haeussler *et al.* 2016) which aligns 20mers using BWA (Li and Durbin 2009) in iterative mode, ranks off-targets by CFD or MIT scores and calculates the Fusi/Doench 2016 efficiency scores (Fusi *et al.* 2015; Doench *et al.* 2016). Efficiency scores were also translated to percentiles for better ease of use, e.g. a raw Fusi/Doench score of 60 translates to the 80th percentile, meaning that 80% of scored guides are lower than 60. All results are then written to a UCSC BigBed file (Raney *et al.* 2014) for interactive visualization. The BigBed file can be loaded into all popular genome browsers, like Ensembl (Yates *et al.* 2016), IGV (Robinson *et al.* 2011) or GBrowse (Stein *et al.* 2002). The track is available on the ci2 assembly on the UCSC Genome Browser (http://genome.ucsc.edu), in the track group “Genes and Gene Predictions”. The source code is available as part of the UCSC Genome Browser source tree at: https://github.com/ucscGenomeBrowser/kent/tree/master/src/hg/makeDb/crisprTrack

### Embryo imaging

Fluorescent *in situ* hybridization of *eGFP* or *Foxf* coupled to immunohistochemsitry was carried out as previously described (Beh *et al.* 2007; Stolfi *et al.* 2014). Images were taken on a Leica Microsystems inverted TCS SP8 X confocal microscope or a Leica DM2500 epifluorescence microscope. Mouse monoclonal anti-β-Gal Z3781 (Promega, Madison, WI) was used diluted at 1:500. Goat anti-Mouse IgG (H+L) Secondary Antibody Alexa Fluor 568 conjugate (Life Technologies) was used diluted at 1:500.

## Acknowledgments

We are grateful to Farhana Salek, Kristyn Millan, and Aakarsha Pandey for technical assistance; Tara Rock for advice on next-generation sequencing; Rahul Satija for sequencing the libraries and for his invaluable insights into the sgRNA sequence analysis; Justin S. Bois, Shyam Saladi, Elena K. Perry, and the High Performance Computing team at NYU for their help troubleshooting the bioinformatic analysis. This work was funded by an NIH K99 HD084814 award to A.S., NIH R01 GM096032 award to L.C., and an NYU Biology Masters Research Grant to S.G.

---

**Supplementary Figure 1.**
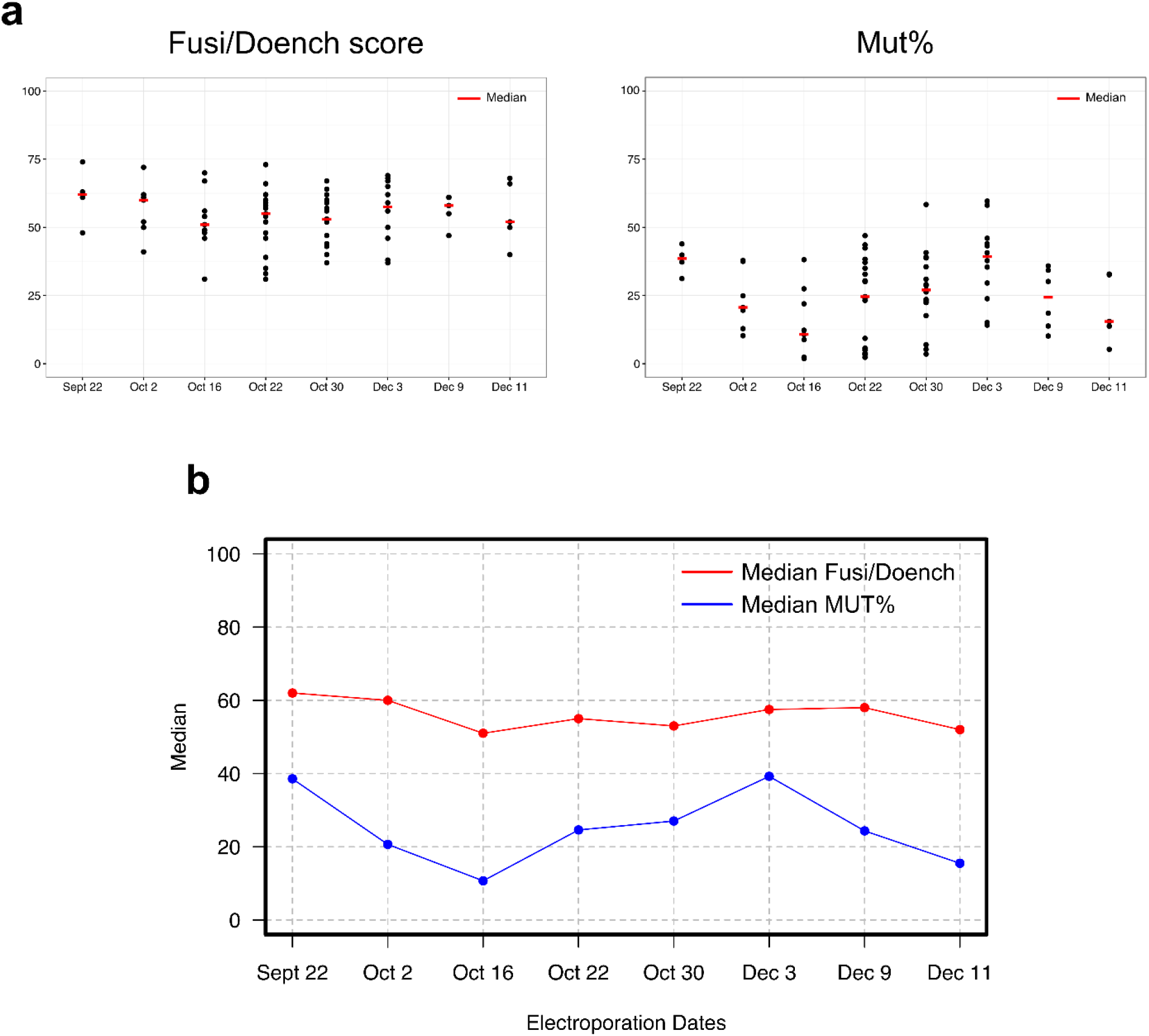
Fusi/Doench scores and mutagenesis efficacies plotted by electroporation date. **a)**Fusi/Doench scores (left) and mutagenesis efficacy estimates (Mut%, right) for individual sgRNAs tested, grouped by electroporation date. **b)** Plot of median values indicated in (a). While variation in Fusi/Doench score within and between dates should be random, mut% could in theory be affected by electroporation efficiency variation, or embryo batch effects. Despite the narrower range of Fusi/Doench scores, trends were similar for both datasets.

**Supplementary Figure 2.**
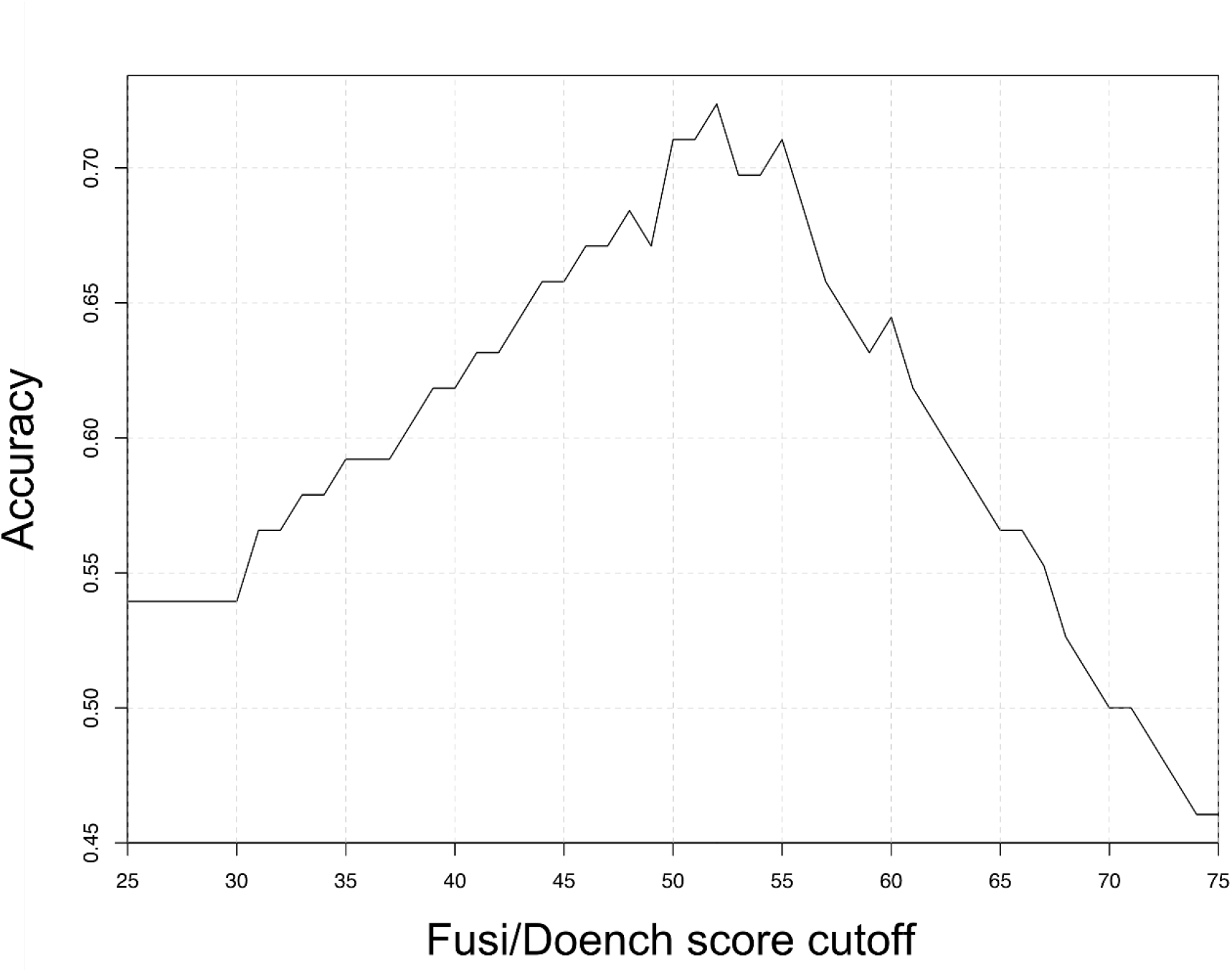
Accuracy of good vs. bad sgRNA classification using different Fusi/Doench score cutoffs. Accuracy was defined as the percentage of correctly classified instances (True Positives + True Negatives)/(True Positives + True Negatives + False Positives + False Negatives). The maximum accuracy was 0.72, using a cutoff of 52. See **Supplementary Table 2** for data.

**Supplementary Figure 3.**
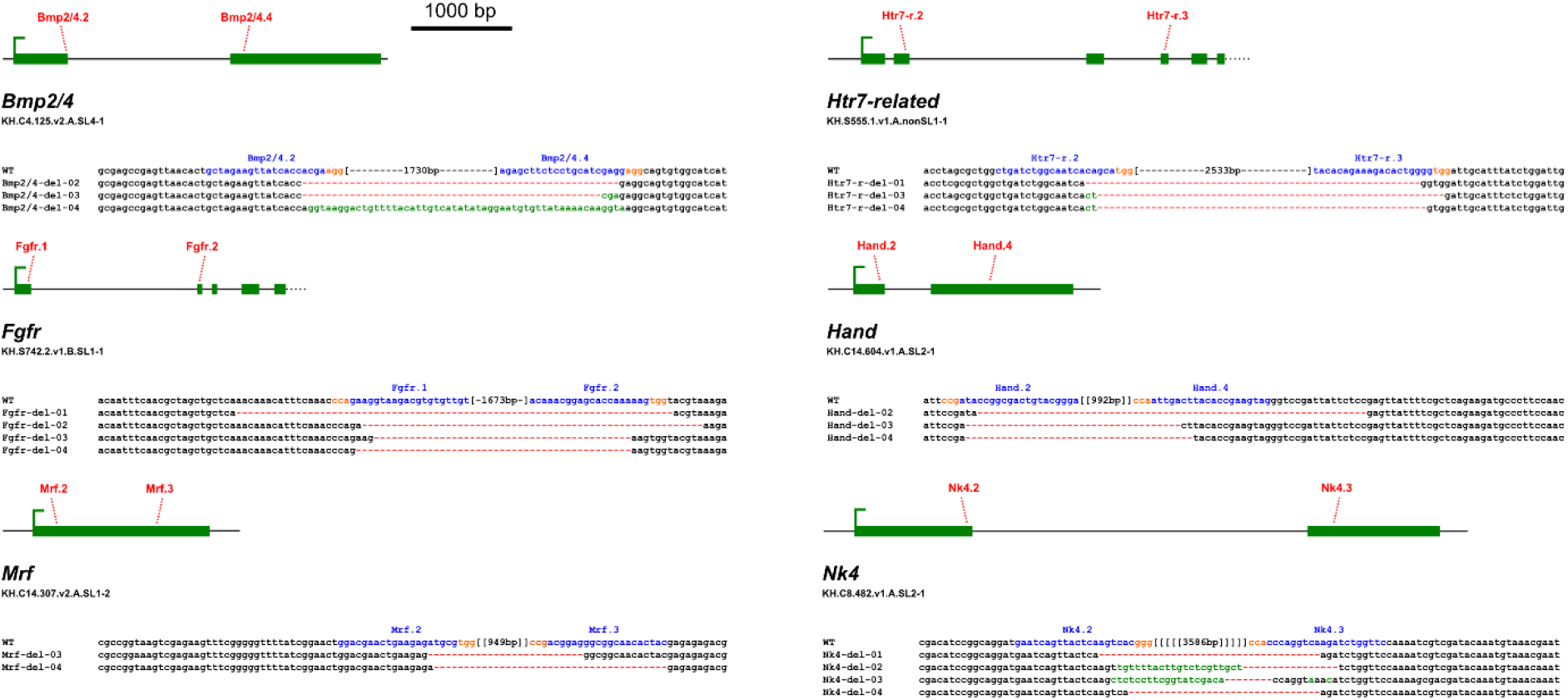
Other examples of large deletions obtained by combinatorial action of two sgRNAs. Sequence alignments of clones for each locus, amplified from embryos in which two sgRNAs were used for CRISPR/Cas9-induced site mutagenesis.

**Supplementary Figure 3.**
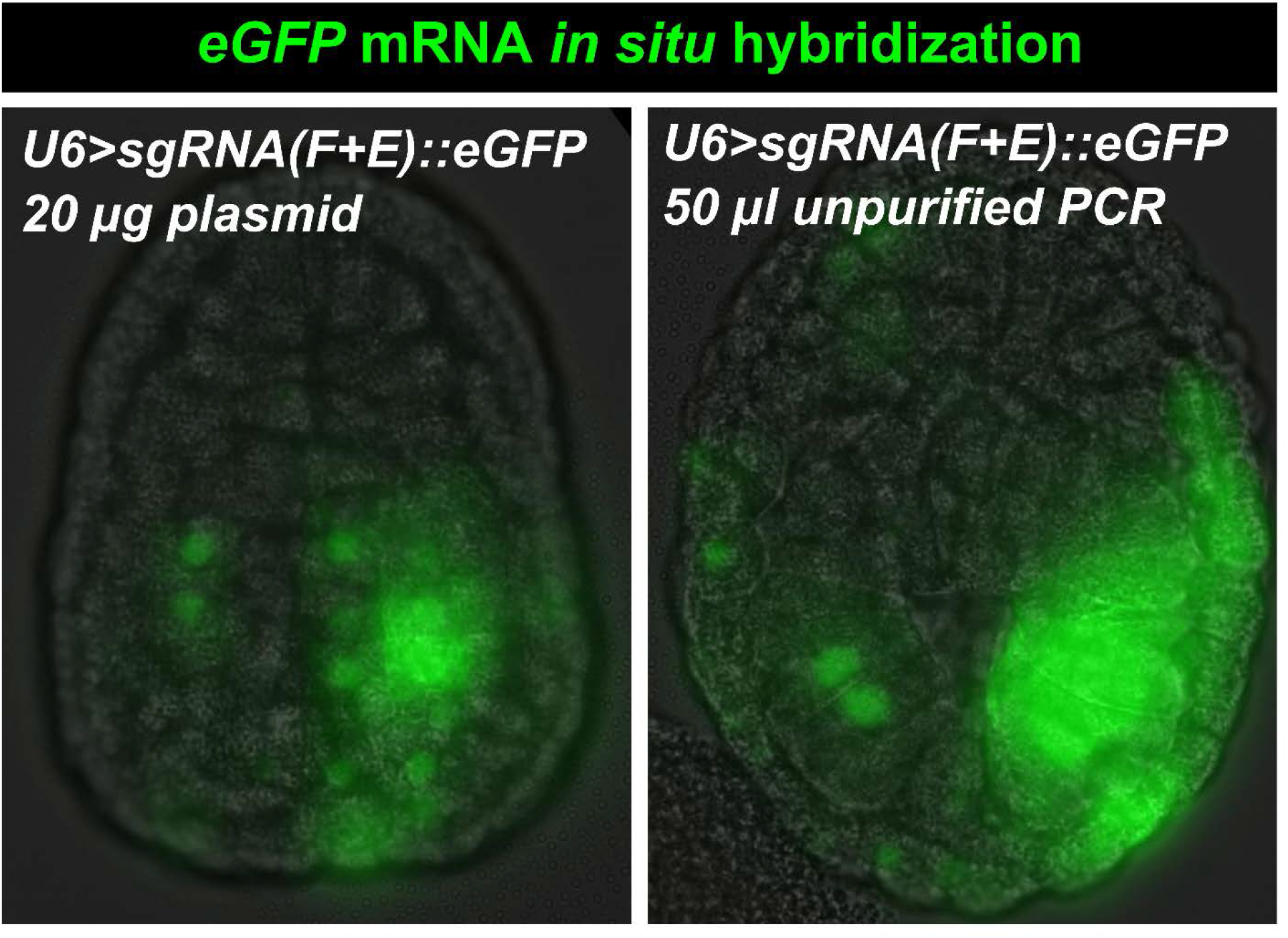
Evidence of *in vivo* transcription from electroporated, unpurified PCR products. In situ hybridization of eGFP in late gastrula/early neural stage embryos electroporated with either *U6*>*sgRNA(F+E)::eGFP* plasmid (20 µg) or unpurified PCR product (50 µl, ˜5 µg DNA).

**Supplementary Figure 5.**
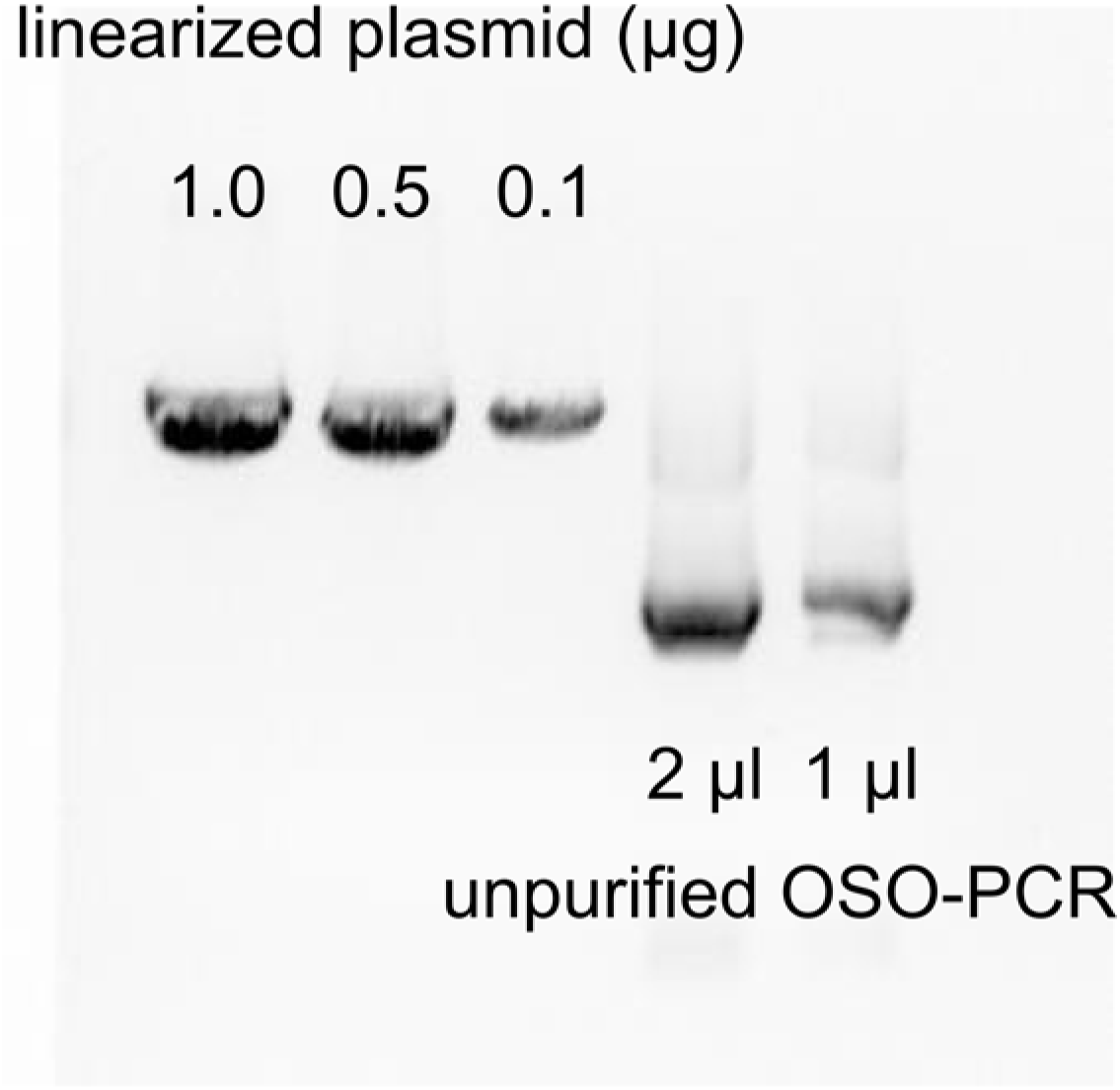
Quantification of OSO-PCR products. Image of gel electrophoresis of varying amounts of linearized plasmid and unpurified OSO-PCR products. Pixel intensity analysis in ImageJ was performed as previously described (Stolfi *et al.* 2014), and indicated that the sgRNA expression cassette in unpurified OSO-PCR reactions are at a concentration of approximately 100 ng/µl.

**Supplementary Figure 6.**
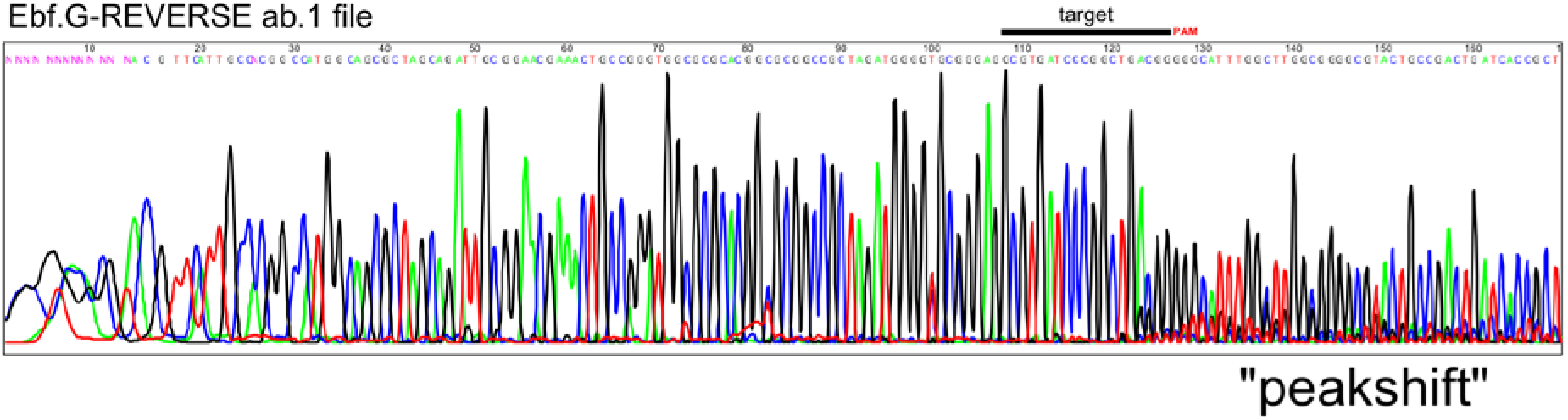
Detection of CRISPR/Cas9-induced indels by Sanger sequencing. Chromatogram of Ebf amplicon from embryos targeted using the Ebf.G sgRNA. “Peakshift” shows superimposed sequence peaks as a result of the resulting mix of mutant alleles bearing short indels around the Ebf.G target site and PAM. This peakshift can be quantified and corrected to produce a precise quantification of CRISPR/Cas9-mediated mutagenesis (see **Materials and methods** for details).

## Supplemental Protocol

### ONE-STEP OVERLAP PCR (OSO-PCR) TO MAKE READY-TO-ELECTROPORATE SINGLE GUIDE RNA (sgRNA) EXPRESSION CASSETTES – *updated 09/21/2016*

> **Companion manuscript:**
>
> Evaluation and rational design of guide RNAs for efficient CRISPR/Cas9-mediated mutagenesis in *Ciona*
>
> Shashank Gandhi, Maximilian Haeussler, Florian Razy-Krajka, Lionel Christiaen, and Alberto Stolfi

Primers for OSO-PCR ready to be ordered can be obtained from the CRISPOR sgRNA prediction and design website (http://crispor.tefor.net), which also checks for known single-nucleotide polymorphisms (SNPs) and potential off-targets in the genome. You can also check for polymorphisms using the Kyoto University Ghost Database genome browser (http://ghost.zool.kyoto-u.ac.jp/cgi-bin/gb2/gbrowse/kh/). You should avoid sgRNAs targeting known SNPs or naturally occurring indels, since Cas9 cutting depends on perfect target sequence match. To design OSO-PCR primers *de novo*, follow the instructions:

1- Select your target, as identified by online tools such as CRISPOR (see above).

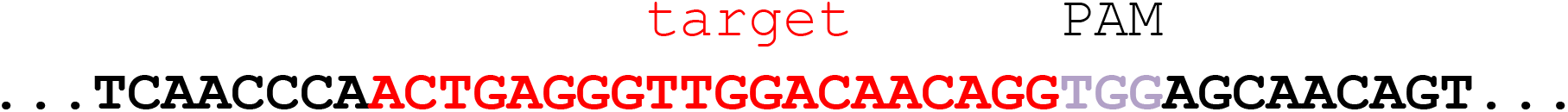

2- A target (the protospacer) is given as N(20). If the target sequence contains too many T’s (three or more T’s clustered together tend to terminate transcription), or if it spans many known naturally-occurring polymorphisms, or has a high number of potential off-targets, discard it.

3- For transcription initiation from U6 promoter, replace the first base of the target with a “G”, to give a G+(N)19 sequence.

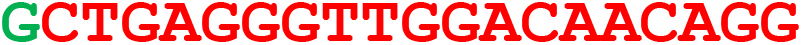

4- Append “GTTTAAGAGCTATGCTGGAAACAG” to the 3’ end of the sequence. This entire sequence is now the forward primer used to PCR the sgRNA scaffold part of the cassette (“OSO forward” primer)

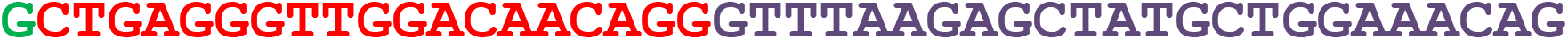

5- Copy reverse complement of G+N(19), append “ATCTATACCATCGGATGCCTTC” to the 3’ end of this. This is now the reverse primer to PCR the U6 promoter part of the cassette (“OSO reverse” primer)

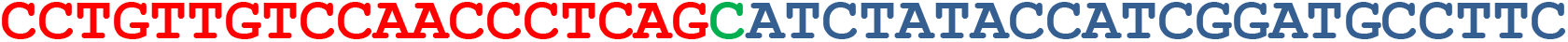

6- Set up a PCR reaction using the following components in the exact amounts described. The amounts/concentrations/proportions are critical for the one-step overlap reaction to occur seamlessly. Also, it is very important to eliminate all sources of contamination, otherwise you may re-amplify sgRNAs already in heavy use in the lab. Template plasmids are available from Addgene (https://www.addgene.org/Lionel_Christiaen/):

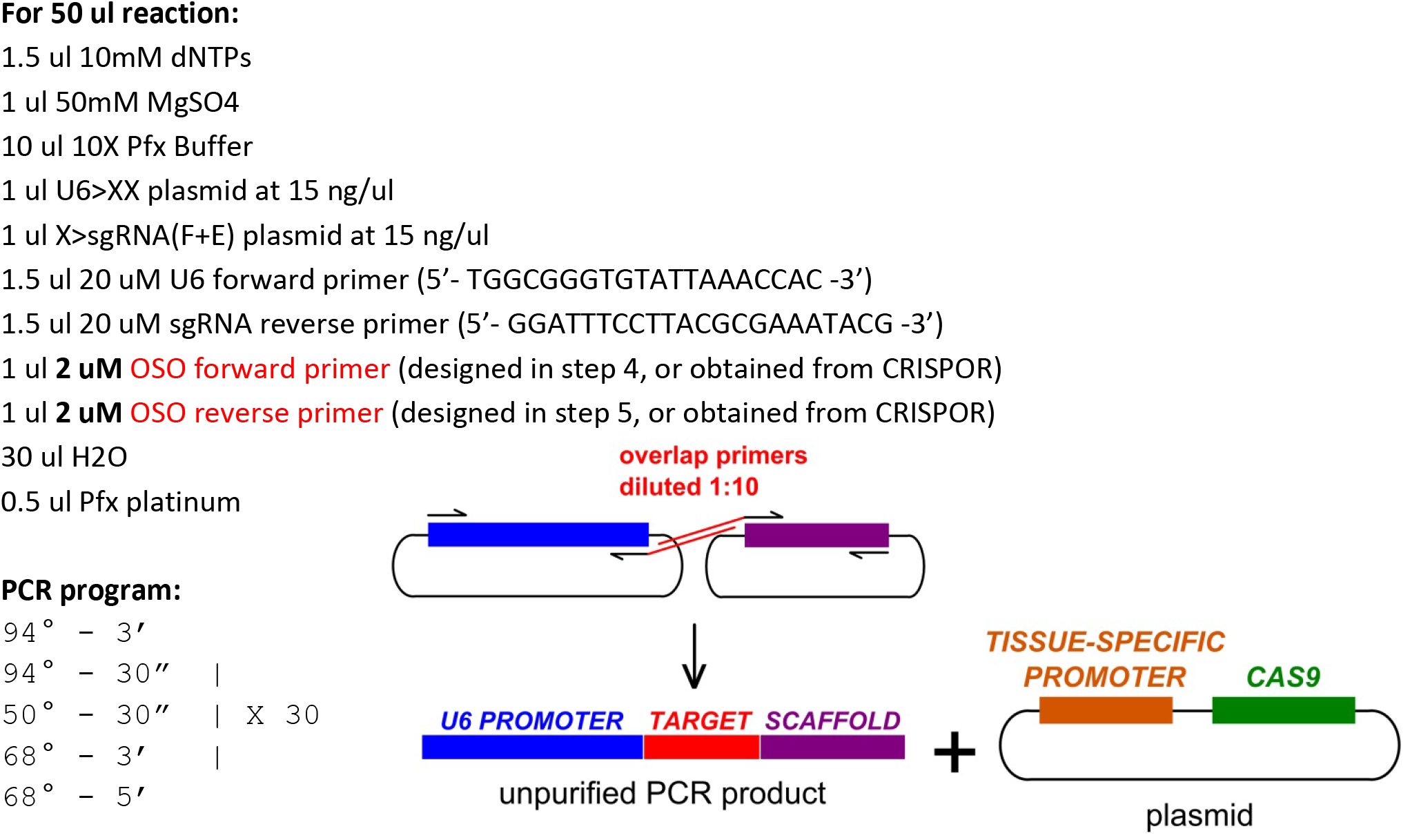

The 1:10 dilution of your custom overlap target-specific primers will force the “fusion” of the entire cassette later in the reaction, when these primers are depleted from the solution through incorporation into the PCR products.

7- Run 2 ul of the PCR reaction on a gel. There should be a strong band at ˜1.2 kbp. If the band is only 1 kbp, the fusion did not occur. The success rate in our hands is ˜94%. If possible, run alongside positive control (PCR on verified sgRNA plasmid template using same primers).

OSO-PCR products can be electroporated as is, un-purified. 25 μl appears to be sufficient to recapitulate effects of sgRNAs delivered by traditional plasmid electroporation, but this volume can be adjusted accordingly. If you need to clone the cassette into a plasmid, you can use the product as template for additional PCRs using the outer primers with added overhangs for restriction enzyme or Clontech In- Fusion cloning.

